# MYC and p53 alterations cooperate through VEGF signaling to repress cytotoxic T cell and immunotherapy responses in prostate cancer

**DOI:** 10.1101/2024.07.24.604943

**Authors:** Katherine C. Murphy, Kelly D. DeMarco, Lin Zhou, Yvette Lopez-Diaz, Yu-jui Ho, Junhui Li, Shi Bai, Karl Simin, Lihua Julie Zhu, Arthur M. Mercurio, Marcus Ruscetti

## Abstract

Patients with castration-resistant prostate cancer (CRPC) are generally unresponsive to tumor targeted and immunotherapies. Whether genetic alterations acquired during the evolution of CRPC impact immune and immunotherapy responses is largely unknown. Using our innovative electroporation-based mouse models, we generated distinct genetic subtypes of CRPC found in patients and uncovered unique immune microenvironments. Specifically, mouse and human prostate tumors with *MYC* amplification and *p53* disruption had weak cytotoxic lymphocyte infiltration and an overall dismal prognosis. MYC and p53 cooperated to induce tumor intrinsic secretion of VEGF, which by signaling through VEGFR2 expressed on CD8^+^ T cells, could directly inhibit T cell activity. Targeting VEGF-VEGFR2 signaling *in vivo* led to CD8^+^ T cell-mediated tumor and metastasis growth suppression and significantly increased overall survival in *MYC* and *p53* altered CPRC. VEGFR2 blockade also led to induction of PD-L1, and in combination with PD-L1 immune checkpoint blockade produced anti-tumor efficacy in multiple preclinical CRPC mouse models. Thus, our results identify a genetic mechanism of immune suppression through VEGF signaling in prostate cancer that can be targeted to reactivate immune and immunotherapy responses in an aggressive subtype of CRPC.

**Significance:** Though immune checkpoint blockade (ICB) therapies can achieve curative responses in many treatment-refractory cancers, they have limited efficacy in CRPC. Here we identify a genetic mechanism by which VEGF contributes to T cell suppression, and demonstrate that VEGFR2 blockade can potentiate the effects of PD-L1 ICB to immunologically treat CRPC.

## INTRODUCTION

Prostate cancer is the leading cancer afflicting American men, with 1 in 8 males diagnosed with prostate cancer in their lifetime (1). The standard-of-care for advanced prostate cancer is a form of androgen-deprivation therapy (ADT), to which most patients initially respond well (2). However, up to 30% of prostate cancers will relapse with castration-resistant prostate cancer (CRPC), which is no longer responsive to hormone therapy and quickly becomes metastatic (3,4). While great strides have been made to develop next generation androgen receptor (AR) signaling inhibitors (ARSIs) that are now clinically approved (5), they generally offer temporary benefit, and metastatic CRPC (mCRPC) remains intractable. An alternative therapeutic avenue in prostate and other treatment-refractory solid tumor malignancies has been the use of immunotherapies to stimulate immune recognition and clearance of local and disseminated tumor cells. Indeed, Sipuleucel-T (Provenge) was the first cancer vaccine to be approved by the FDA and achieved designation in the setting of mCRPC (6). Still, its effects on overall survival remain marginal at-best for patients (7,8), and other immunotherapy modalities such as anti-CTLA-4 and PD-1/PD-L1 immune checkpoint blockade (ICB) that have been curative in other cancers are generally ineffective in prostate malignancies (9–11). This lack of durable immunotherapy responses is believed to be due to the inherently “cold” tumor microenvironment (TME) of prostate cancer that is devoid of the cytotoxic lymphocytes and enriched in suppressive myeloid cell populations (12). Thus, it will be critical to understand the mechanisms contributing to the immune suppressive prostate TME in order to design more effective immunotherapy strategies for CRPC.

Large-scale analyzes of patient samples have revealed genomic, molecular, and histological subtype classifications of CRPC (13–20). Though the majority of CRPCs remain AR-dependent through AR amplification or splice variants (21,22), many become an AR-independent form of aggressive variant prostate cancer (AVPC) through acquisition of additional genetic or epigenetic alterations or lineage conversion into neuroendocrine prostate cancer (NEPC) (23–25). Such genetic alterations acquired in CRPC include amplification of oncogenic *MYC* and *MYCN*, mutations or deletions in tumor suppressor genes (TSGs) such as *TP53*, *PTEN*, *RB1*, and *APC,* and perturbations in DNA repair pathways (15,16). Interestingly, recent work has demonstrated that, albeit rare, CRPCs harboring alterations in DNA damage (*CDK12*) and mismatch repair (*MSH2, MLH1*) genes resulting in microsatellite instability present with an inflamed TME with increased antigen presentation and better response rates to anti-PD-1 ICB (14,26–28). In contrast, previous studies in prostate and other cancer types revealed a role for more prevalent genetic alterations such as *MYC* induction and *TP53*, *PTEN*, and *APC* inactivation in promoting the infiltration of myeloid-derived suppressor cells (MDSCs) and macrophages, reducing antigen presentation by tumor cells and dendritic cells (DCs), and suppressing interferon signaling necessary for both innate and adaptive immunity (29–37). As such, understanding how genetic alterations that frequently co-occur in prostate cancer impact the immune landscape could lead not only to better stratification of patients for precision medicine, but also to new therapeutic approaches to treat different subtypes of CRPC.

To model the complex and compound genetic alterations commonly associated with AVPC in a rapid and flexible manner, we previously developed electroporation-based non-germline genetically engineered mouse models (EPO-GEMMs), whereby oncogenes can be expressed by transposon-mediated transgenesis and TSGs inactivated by CRISPR/Cas9-mediated genomic editing to generate prostate tumors *de novo* and *in situ* in adult animals (38,39). These models recapitulate the histological and molecular phenotypes associated with human AVPC, and display low AR expression and indifference to castration indicative of CRPC. Given this platform can be used to generate prostate tumors in their resident TME with an intact immune system, we generated a suite of genetically-defined EPO-GEMMs to identify the unique immune landscapes of different genetic subtypes and explore tumor intrinsic mechanisms of immune suppression. In doing so, we uncovered a novel mechanism of immune suppression in a lethal AVPC subtype driven by *MYC* and *Tp53* (hereafter *p53*) co-alterations that presents an actionable target to remodel the “cold” prostate TME and potentiate ICB responses in CRPC.

## RESULTS

### Prostate cancer EPO-GEMMs exhibit genotype-specific differences in immune landscape

We previously developed electroporation-based non-germline genetically engineered mouse models (EPO-GEMMs) harboring transposition-mediated human c-MYC (MYC) overexpression and CRISPR/Cas9-mediated *Pten* or *p53* disruption that produce lethal and metastatic prostate cancer *de novo* and *in situ* in the anterior lobe of the prostate of adult mice with high penetrance (83% and 76%, respectively) (Fig. 1A-B) (38). Given the rapid nature and flexibility to engineer various compound oncogene and TSG alterations in these models, we used this platform to explore the impact of different genetic alterations commonly associated with human CRPC on the immune landscape (Fig. 1C). In addition to MYC-driven prostate cancer EPO-GEMMs harboring compound human *MYC* overexpression with *Pten* (hereafter *MPten*) or *p53* (hereafter *MP*) inactivation we previously characterized (38), we also generated a new EPO-GEMM model defined by CRISPR-mediated disruption of three TSGs, *Pten*, *p53*, and *Rb1* (hereafter *PtPRb*), in the absence of *MYC* alterations (Fig. 1A; Supplementary Fig. S1A,B). *PtPRb* EPO-GEMMs developed lethal prostate cancer with a high penetrance (83%) but with a longer latency (median survival of 167 days) compared to MYC-driven *MPten* and *MP* genetic subtypes (median survival 74 and 90 days, respectively) (Fig. 1B). Like *MP* and *MPten* tumors, *PtPRb* tumors had low expression of the androgen receptor (AR) and the luminal marker CK8, and were unresponsive to ADT, indicative of a poorly differentiated and aggressive variant prostate cancer (AVPC) likely to be castration-resistant (20,25) (Supplementary Fig. S1C-D). Consistent with the association of these co-alterations with small cell or neuroendocrine (NE) differentiation in human CRPC (23,40,41), *PtPRb* tumors also expressed neuroendocrine markers Synaptophysin (SYP), NEUROD1, and ASCL1, as well as a small cell morphology suggestive of a NE phenotype (Supplementary Fig. S1D).

**Figure 1.**
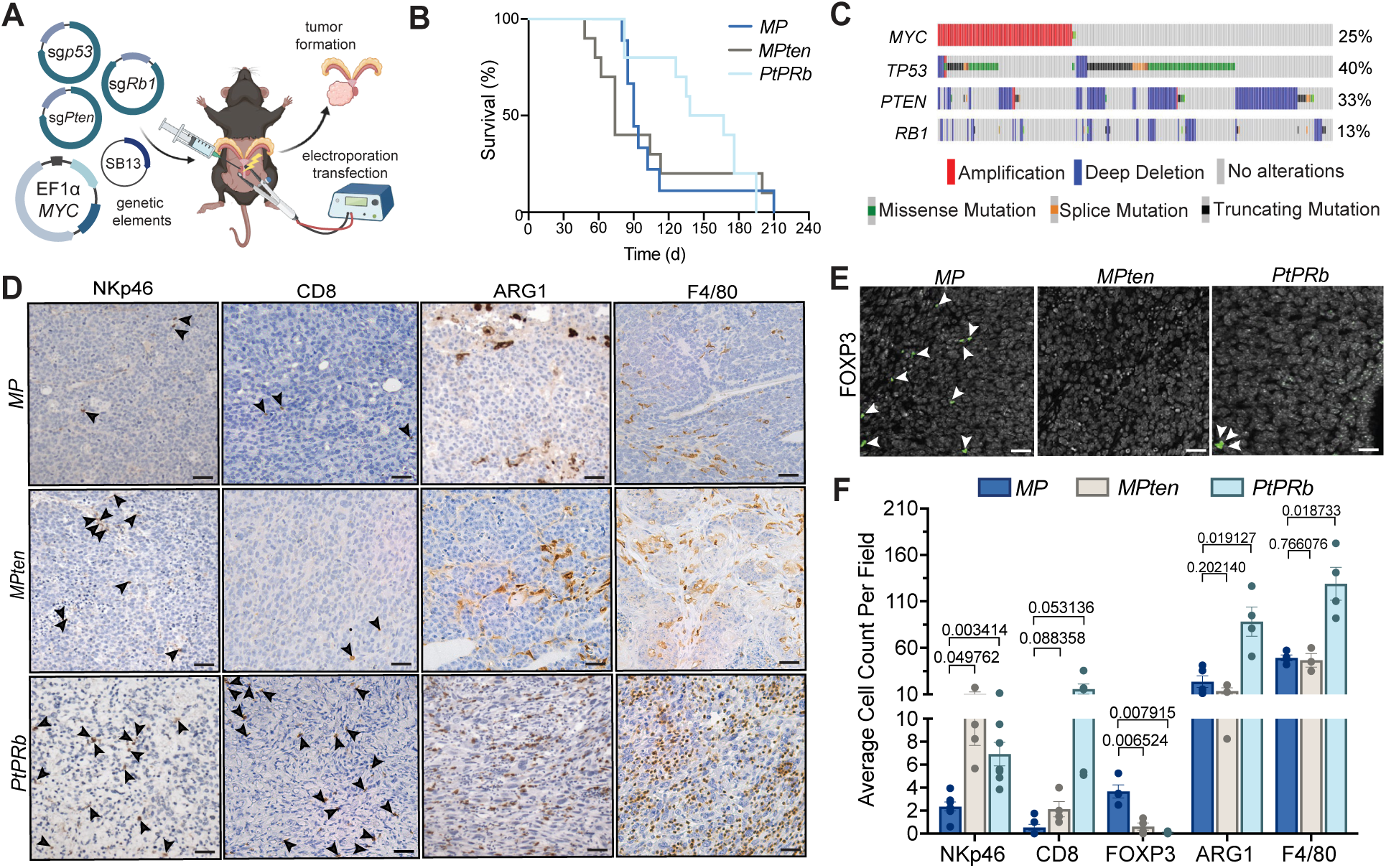
EPO-GEMM prostate cancer models reveal genetically-defined changes in the immune TME. **A,** Schematic of electroporation-based non-germline genetically engineered mouse model (EPO-GEMM) generation and specific oncogene and tumor suppressor gene (TSG) alterations engineered. Created with Biorender.com. SB, sleeping beauty transposase. Sg, single guide RNA. **B**, Kaplan-Meier survival curves of EPO-GEMMs produced in C57BL/6 mice harboring prostate tumors with indicated genotypes (n = 9-11 mice per group). *MP*, *MYC;p53^-/-^*. *MPten*, *MYC;Pten^-/-^*. *PtPRb*, *Pten^-/-^;p53^-/-^;Rb1^-/-^*. **C,** OncoPrint displaying types and frequencies of genomic alterations in *MYC*, *TP53*, *PTEN*, and *RB1* in metastatic CRPC (mCRPC) patient samples from Stand Up to Cancer (SU2C) datasets (15) (n = 444 patient samples). Generated on cBioPortal.org. **D-E,** Representative immunohistochemical (IHC) (**D**) and immunofluorescence (IF) (**E**) staining of *MP*, *MPten*, and *PtPRb* EPO-GEMM prostate tumors harvested at endpoint. Arrowheads indicate positive staining for immune cells. Scale bars, 50μm. **F,** Quantification of NKp46^+^ NK cells, CD8^+^ T cells, FOXP3^+^ regulatory T cells (Tregs), Arginase1^+^ (ARG1) suppressive myeloid cells, and F4/80^+^ macrophages per field (n = 3-7 mice per group). Data represent mean ± SEM. P-values were calculated by two-tailed, unpaired Student’s t-test.

To determine the contribution of MYC and TSG alterations to the immune suppressive prostate cancer landscape, genetically-defined tumors of similar size harvested from these animals at survival endpoint were subjected to immunophenotyping by immunohistochemistry (IHC) or immunofluorescence (IF) analysis. Despite both *MPten* and *MP* prostate tumors having the same driver oncogene, they had distinct immune infiltrates defined by their respective TSG alterations. Whereas both *MPten* and *MP* prostate tumors had accumulation of F4/80^+^ macrophages and suppressive myeloid cells expressing Arginase 1 (ARG1), *MP* tumors had reduced numbers of cytotoxic Natural Killer (NK) and CD8^+^ T cells and an increase in regulatory T cells (T_regs_) compared to *MPten* tumors (Fig. 1D-F). Strikingly, *PtPRb* tumors formed in the absence of MYC induction displayed a significant increase in both cytotoxic NK and CD8^+^ T lymphocytes, as well as ARG1^+^ and F4/80^+^ myeloid cells compared to *MP* subtypes also harboring *p53* alterations (Fig. 1D-F). These genotype-specific immune phenotypes were further validated in prostate tumors from mice transplanted with *MP*, *MPten,* and *PtPRb* cell lines generated from EPO-GEMM tumors, with *PtPRb* tumors having the largest immune infiltrate and *MP* tumors displaying reduced numbers of lymphoid and myeloid cells in comparison as assessed by IHC analysis (Supplementary Fig. S1E-F). Together, these findings demonstrate that *MYC*, and to an even greater extent compound *MYC* activation and *p53* loss, lead to cytotoxic lymphocyte suppression in prostate cancer.

### Compound *MYC* and *p53* alterations are associated with poor outcomes and immune suppression in human CRPC

To evaluate the clinical impact of *MYC* alterations on immune suppression in human prostate cancer, we first stained primary, surgically resected prostate cancer tumor samples of various Gleason Scores we obtained from the UMass Center for Clinical and Translational Science Biorepository for MYC and markers of NK and CD8^+^ T cells. Consistent with findings in our EPO-GEMM models, human tumors with high MYC expression had significantly fewer CD8^+^ T cell and NK cell infiltrates compared to tumors with low or absent MYC expression as scored by a clinical pathologist (Fig. 2A-B). To more comprehensively investigate the immune landscape of patients with *MYC*, *P53*, and/or *PTEN* alterations, we analyzed expression of immune-related gene signatures within sequenced prostate tumors from The Cancer Genome Atlas (TCGA) (42). Whereas patients with *MYC* alterations alone had no differences in the magnitude of immune infiltration [MImmScore (43)] as well as specific CD8^+^ T cell (44) and NK cell (45) gene signature expression, those with compound *MYC* and *P53* alterations had significantly decreased expression of immune and NK cell signatures, and trended toward reduced expression of CD8^+^ T cell-specific transcripts (Fig. 2C). This decrease in immune-related transcripts was not observed in the context of *PTEN* alterations (Supplementary Fig. S2A). In addition, analysis of a publicly available Stand Up to Cancer (SU2C) dataset (15) revealed that metastatic CRPC (mCRPC) patients harboring *MYC* and *P53* co-alterations had significantly worse overall survival compared to those with *MYC* amplification or *P53* mutation/deletion alone that was not observed with compound *MYC* and *PTEN* alterations (Fig. 2D; Supplementary Fig. 2B), substantiating that this genetic subtype marks an aggressive form of CRPC. These data further support our findings in animal models that *MYC* induction in combination specifically with *p53* disruption, which is found in ∼8-9% of patients (Fig. 1C), leads to an aggressive and immune suppressed subtype of CRPC.

**Figure 2.**
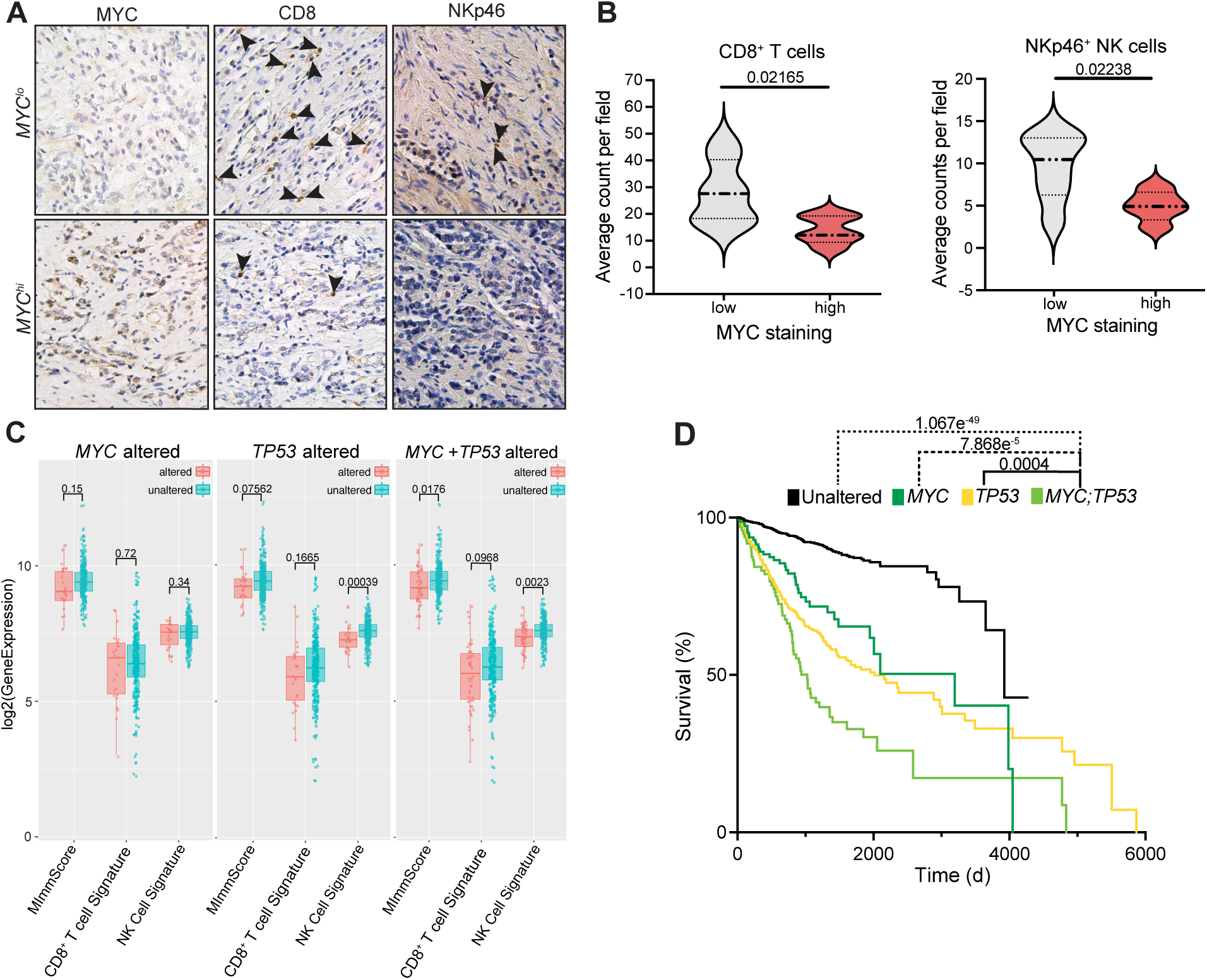
*MYC* and *p53* co-alterations result in immune suppression and more aggressive disease in human prostate cancer. **A,** Representative IHC staining in surgically resected primary prostate cancer patient samples stratified by MYC staining score into high and low groups. Arrowheads indicate positive staining for immune cells. **B,** Quantification of CD8^+^ T cells and NKp46^+^ NK cells in prostate cancer patient samples stratified by MYC staining score into high and low groups (n = 5-7 samples per group). **C,** Box and whisker plots showing gene expression analysis of magnitude of immune infiltration [MImmScore (43)], NK cell (45), and CD8^+^ T cell (44) signatures in primary prostate cancer patient samples from The Cancer Genome Atlas (TCGA) (42) stratified by alterations in *MYC*, *TP53*, or compound *MYC;TP53* (n = 46-235 samples per group). The central line represents the median, the ends of the box the upper and lower quartiles, and whiskers extend to the highest and lowest observations. **D,** Kaplan-Meier survival curves of metastatic CRPC patients harboring prostate tumors with *MYC* or *TP53* alterations alone or in combination from SU2C datasets (15) (n = 38-143 samples per group). Data represent mean ± SEM. P-values were calculated by two-tailed, unpaired Student’s t-test (**B**), Wilcoxon test (**C**), and log-rank test (**D**).

### *MYC* and *p53* alterations combine to repress inflammatory signaling and induce VEGF secretion

We next wanted to determine the tumor cell intrinsic mechanisms by which *MYC* overexpression and *p53* loss cooperate to mediate immune suppression. RNA sequencing (RNA-seq) analysis of bulk prostate tumor samples from genetically defined EPO-GEMM animals revealed that both *MP* and *MPten* tumors had reduced expression of antigen presentation and processing genes (*B2m, H2-d1, H2-k1, Tap1/2, Erap1*) necessary for effective antigen-dependent T cell responses, as well as stimulatory ligands necessary for NK cell engagement (*Ulbp1*, *H60b/c*, *Raet1d/e*) as compared to *PtPRb* tumors lacking *MYC* overexpression (Fig. 3A). *MP* and *MPten* cell lines propagated from EPO-GEMM tumors also had significantly reduced major histocompatibility complex (MHC) Class I (MHC-I) surface levels in comparison to *PtPRb* lines (Supplementary Fig. S3A-B). Gene Set Enrichment Analysis (GSEA) of RNA-seq data from primary tumors demonstrated significant enrichment of genes related to inflammatory, NF-κB, and Type I interferon signaling almost exclusively in *PtPRb* as compared *MP and MPten* tumors (Fig. 3B; Supplementary Fig. 3C). Interestingly, when comparing between *MYC*-driven subtypes, we observed reduced enrichment of these inflammatory response gene sets in *MP* compared to *MPten* subtypes (Fig. 3B), indicating that *p53* TSG loss in the context of *MYC* induction may further suppress inflammatory pathways in prostate tumors that could contribute to an immune suppressed TME.

**Figure 3.**
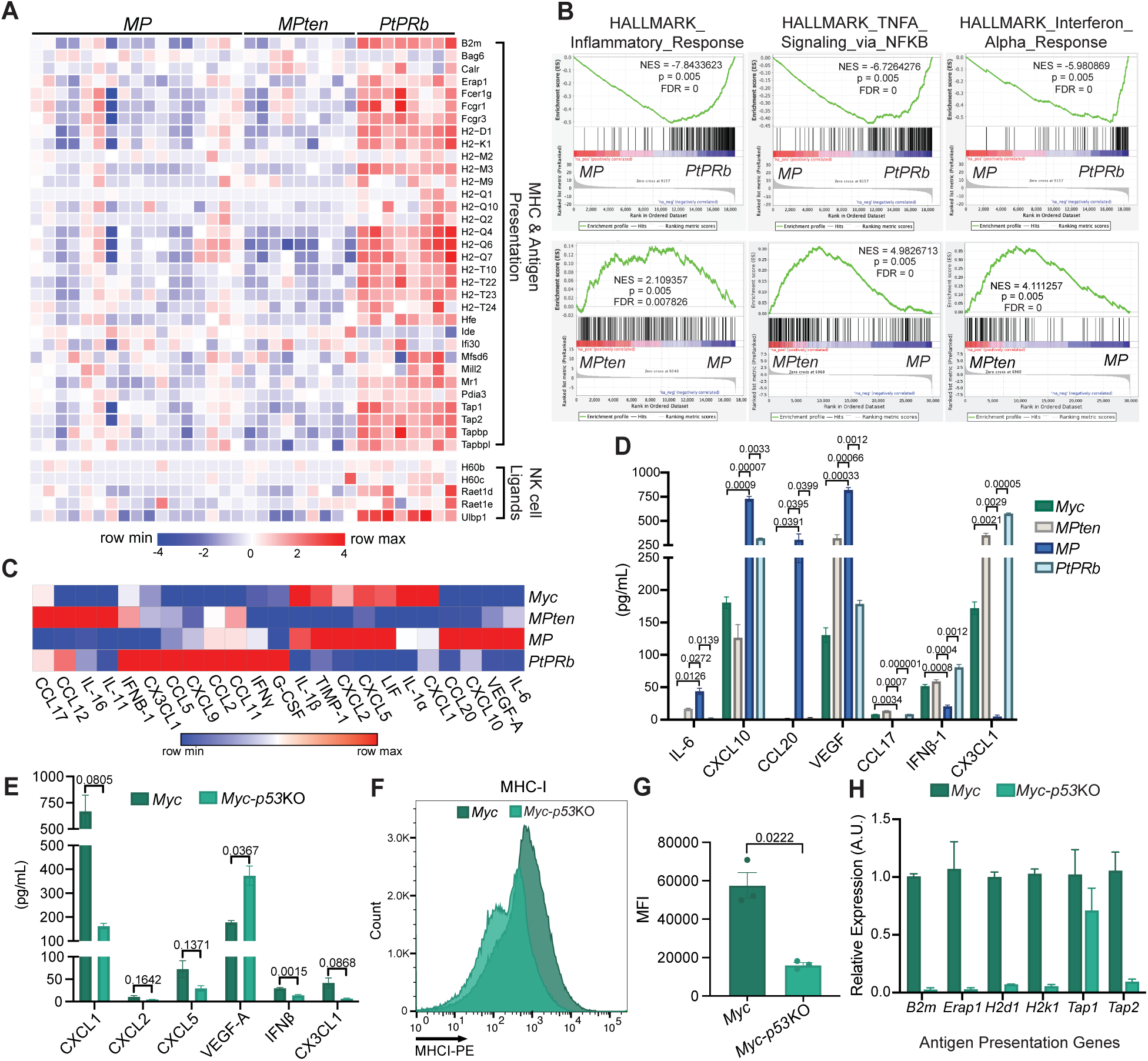
*MYC* induction and *p53* disruption cooperate to repress inflammatory signaling and stimulate VEGF secretion from prostate tumor cells. **A,** Heatmap of major histocompatibility complex (MHC), antigen presentation, and NK cell ligand gene expression in *MP, MPten,* and *PtPRb* EPO-GEMM tumors from bulk RNA-seq analysis (n = 8-17 mice per group). **B,** Gene Set Enrichment Analysis (GSEA) of inflammatory, NFκB, and interferon (IFN) signaling gene sets in indicated EPO-GEMM tumors (n = 8-17 mice per group). NES, normalized Enrichment Score. **C,** Heatmap of cytokine array analysis results from *MycCaP* (*Myc*) cells and *MPten*, *MP*, and *PtPRb* EPO-GEMM-derived cell lines. Data is representative of mean of 3 biological replicates. **D,** Quantification of protein levels of factors differentially secreted in *MP* compared to other cell lines from cytokine array analysis in (**C)** (n = 3 biological replicates per cell line). **E,** Cytokine array analysis of differentially secreted proteins in parental *Myc-CaP (Myc)* cells compared to those with CRISPR-mediated *p53* knockout (*Myc-p53KO*) (n = 3 biological replicates per cell line). **F-G,** Representative histograms (**F**) and quantification (**G**) of mean fluorescent intensity (MFI) of MHC-I (H-2Kb) expression on *Myc-CaP* (*Myc*) and *Myc-p53*KO tumor cells (n = 3 biological replicates per group). **H,** RT-qPCR analysis of antigen presentation genes in *Myc-CaP (Myc)* and *Myc-p53*KO cells (n = 2 biological replicates associated with 3 technical replicates per group). A.U., arbitrary units. Data represent mean ± SEM. P-values were calculated by two-tailed, unpaired Student’s t-test.

To assess differences in the secretory profile of prostate tumors across MYC-driven genotypes, we performed cytokine array analysis on EPO-GEMM-derived *MP*, *MPten*, and *PtPRb* tumor cell lines, as well as the previously characterized *Myc-CaP* cell line propagated from Hi-Myc mice harboring overexpression of a human *MYC* transgene downstream of androgen response elements (hereafter referred to as *Myc* for simplicity) (46,47). Consistent with our RNA-seq analysis, we observed genotype-specific differences in inflammatory chemokine and cytokine secretion, particularly in the *MP* subtype (Fig. 3C). Whereas quantities of some secreted factors, such as IFNβ, CX3CL1, and CCL17, were significantly decreased in *MP* tumor cells, a number of other proteins, including cytokines IL-6 and chemokines CXCL10 and CCL20, were preferentially induced in the *MP* setting as compared to either *Myc*, *MPten*, or even *PtPRb* cells (Fig. 3C-D). Interestingly, one of the most highly induced factors in *MP* as compared to other prostate tumor cell lines was VEGF-A (hereafter VEGF), which through binding to its canonical receptor VEGFR2 can have a pleiotropic effects on diverse immune and stromal cell types in the TME (48–50).

To further dissect the contribution of p53 loss specifically to changes in inflammatory signaling, we used CRISPR/Cas9 to knockout *p53* in *Myc-CaP* prostate tumor cells (hereafter *Myc-p53*KO) (Supplementary Fig. 3D). When transplanted orthotopically into FVB mice, *Myc*-*p53*KO cell line-derived prostate tumors had decreased CD8^+^ T cell accumulation in the TME compared to parental *Myc-CaP*-derived tumors (Supplementary Fig. S3E), recapitulating the limited CD8^+^ T cell infiltration found in *MP* EPO-GEMM tumors with the same genotype (see Fig. 1D and F). *In vitro*, *Myc*-*p53*KO tumor cells had reduced secreted protein of IFNβ and chemokines such as CXCL1, CXCL2, CXCL5, and CX3CL1 important for both myeloid and lymphoid cell chemotaxis into the TME compared to parental *Myc-CaP* cells (Fig. 3E). *P53* deletion in *Myc-CaP* cells also led to reduced expression MHC-I and antigen presentation/processing genes (Fig. 3F-H). Consistent with *MP* EPO-GEMM lines, the most differentially upregulated secreted factor in *Myc*-*p53*KO lines was VEGF (Fig. 3E). Overall, these data demonstrate that *MYC* overexpression and *p53* loss of function cooperate in a tumor intrinsic manner to not only inhibit inflammatory signaling important for attracting both lymphocytes and myeloid cells and presenting antigen to T cells, but also produce VEGF that could have dynamic effects on the TME of CRPC.

### VEGF signaling directly suppresses CD8^+^ T cells in human and murine prostate cancers

Given our results showing increased VEGF expression in *MP* prostate cancer cell lines, we hypothesized that VEGF signaling may contribute indirectly or directly to immune suppression in this aggressive subtype. Consistent with our *in vitro* findings, VEGF protein levels were significantly higher in *MP* EPO-GEMM primary tumors and *Myc-p53*KO cell line transplant tumors compared to tumors of other genetic subtypes as assessed by IHC analysis (Fig. 4A-B). Co-immunofluorescence (co-IF) analysis demonstrated that VEGF expression in these prostate cancer models was predominantly localized within MYC^+^ tumor cells as opposed to macrophages that are also known to secrete VEGF in the TME, confirming tumor intrinsic VEGF production (Supplementary Fig. S4A). Tumor-derived VEGF signaling was also associated with increased numbers of CD31^+^ blood vessels in *Myc* and *p53* co-altered tumors (Fig. 4C), consistent with the canonical role of VEGF in angiogenesis. However, blood vessels in tumors of the *MP* genetic subtype were smaller and lacked visible open lumens, indicative of reduced vascular integrity that could contribute indirectly to poor extravasation of immune cells into the TME (51) (Fig. 4A; Supplementary Fig. S4B).

**Figure 4.**
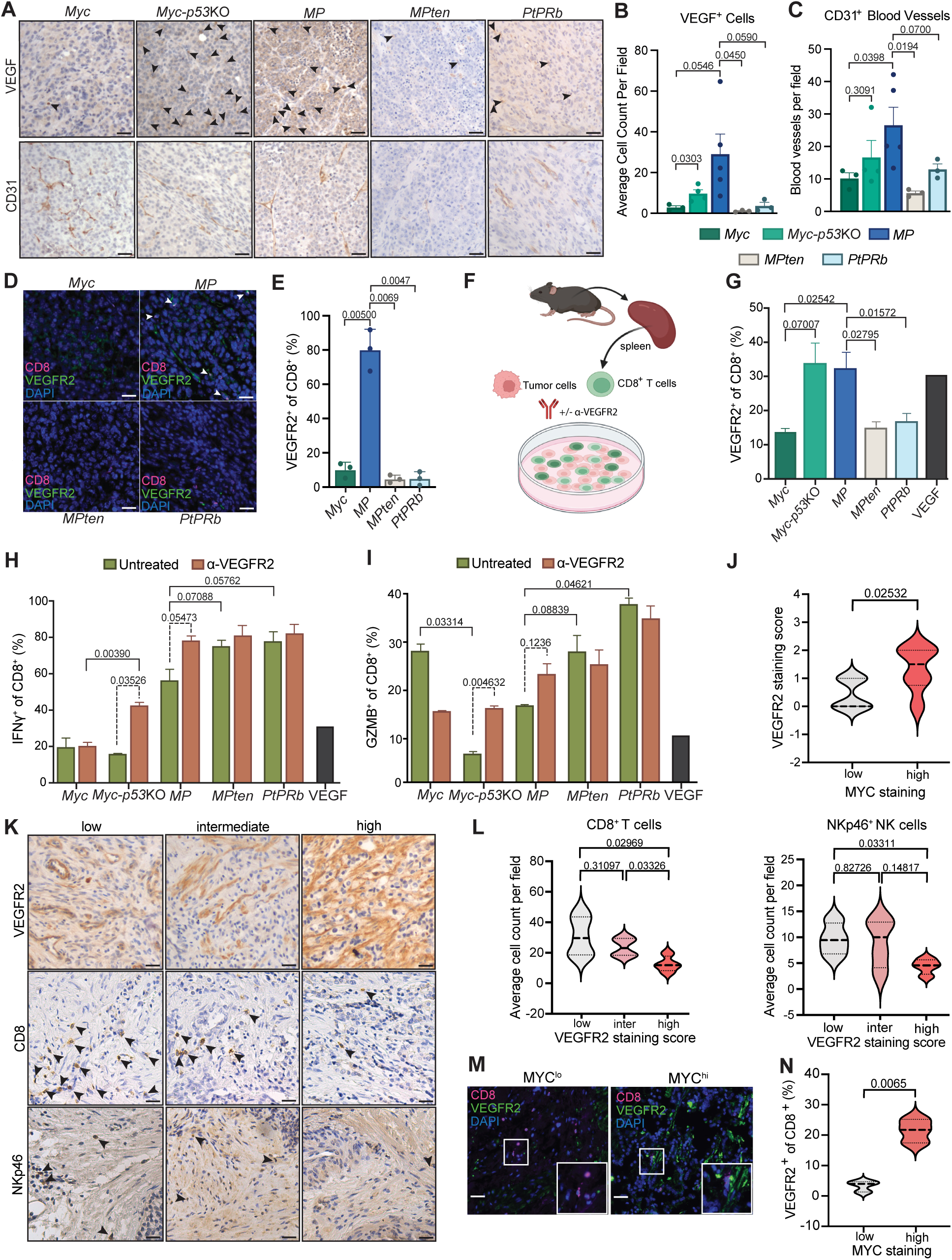
VEGF leads to suppression of VEGFR2-expressing CD8^+^ T cells in murine and human prostate cancer. **A,** Representative IHC staining of prostate tumors from FVB mice transplanted orthotopically with *Myc-CaP (Myc)* or *Myc-p53*KO cells or from C57BL/6 mice transplanted orthotopically with *MP*, *MPten* or *PtPRb* EPO-GEMM-derived cell lines. Arrowheads indicate VEGF positive cells in tumor areas. Scale bars, 50μm. **B-C,** Quantification of VEGF^+^ cells (**B**) and CD31^+^ blood vessels (**C**) per field in **A** (n = 3-4 mice per group). **D,** Representative co-IF staining for CD8 and VEGFR2 in indicated *Myc-CaP* or EPO-GEMM cell line-derived transplant prostate tumors. White arrowheads indicate VEGFR2^+^CD8^+^ double positive cells. Scale bars, 50μm. **E,** Quantification of percentage of CD8^+^ T cells that are VEGFR2^+^from co-IF analysis in **D** (n = 3 mice per group). **F,** Schematic of *ex vivo* tumor-immune co-culture assay using spleen-derived CD8^+^ T cells and murine prostate cancer cell lines. Created with Biorender.com. **G,** Flow cytometry analysis of VEGFR2 expression on CD8^+^ T cells cultured *ex vivo* with indicated prostate cancer cell lines (n = 3 biological replicates per group) or in the presence of recombinant VEGF (50ng/mL). **H-I,** Flow cytometry analysis of IFNγ (**H**) and Granzyme B (GZMB) (**I**) expression in CD8^+^ T cells cultured with indicated prostate cancer cell lines in the presence or absence of a VEGFR2 blocking antibody (DC101; 1µg/mL) (n = 3 biological replicates per group) or recombinant VEGF (50ng/mL). **J,** Quantification of VEGFR2 staining scores in primary prostate cancer patient samples stratified by MYC staining score into high and low groups (n = 7 samples per group). **K,** Representative IHC staining in primary prostate cancer patient samples stratified by VEGFR2 staining score into low, intermediate, and high groups. Arrowheads indicate positive staining for immune cells. Scale bars, 50μm. **L,** Quantification of CD8^+^ T cell and NKp46^+^ NK cell numbers in primary prostate cancer patient samples stratified by VEGFR2 staining score in **K** (n = 4-6 samples per group). **M,** Representative co-IF staining of CD8 and VEGFR2 expression in primary prostate cancer patient tumors stratified by MYC staining score into high and low groups. Scale bars, 50μm. **N,** Quantification of percentage of CD8^+^ T cells that are VEGFR2^+^ in MYC^hi^ and MYC^lo^ patient prostate tumors in **M** (n = 5-7 per group). Data represent mean ± SEM. P-values were calculated by two-tailed, unpaired Student’s t-test.

Recent evidence suggests that VEGF can also directly impact T cell phenotypes in cancer (52–54). Indeed, co-IF analysis revealed that the majority of CD8^+^ T cells in *MP* tumors, but not in other genetic tumor contexts, expressed the VEGF receptor VEGFR2 (Fig. 4D-E). This prompted us to investigate whether VEGF produced by *MP* prostate tumor cells could have a direct effect on the function of CD8^+^ T cell expressing VEGFR2. To this end, we performed *ex vivo* tumor-immune co-culture assays with prostate cancer cells and CD8^+^ T cells isolated from spleens of wild-type (WT) FVB male mice (Fig. 4F). Interestingly, we observed increased expression of VEGFR2 on CD8^+^ T cells co-cultured with *MP* and *Myc-p53*KO tumor cells compared to other genetically-defined prostate cancer lines and to a similar degree as T cells directly stimulated with recombinant VEGF as measured by flow cytometry analysis (Fig. 4G; Supplementary Fig. S4C). Moreover, CD8^+^ T cells co-cultured with *MYC and p53* co-altered tumor cells were less activated than those cultured with *MPten* and *PtPRb* cell lines as assessed by IFNγ and GZMB expression (Fig. 4H-I). Remarkably, while the addition of a VEGFR2 blocking antibody (α-VEGFR2; DC101) to the co-cultures containing *Myc*, *MPten*, and *PtPRb* cells had no impact on T cell activation, VEGFR2 blockade restored both IFNγ and GZMB expression in CD8^+^ T cells exposed to *MP* and *Myc-p53*KO tumor cells to similar levels as those cultured with other genetic prostate cancer subtypes (Fig. 4H-I). These results suggest a direct functional role for VEGF in cytotoxic T cell suppression specifically in *MP* altered prostate cancer.

Analysis of prostate cancer patient samples confirmed these findings from our murine prostate cancer models. First, IHC analysis demonstrated a significantly higher VEGFR2 staining score in prostate cancer patient samples with high MYC expression (Fig. 4J; Supplementary Fig. S4D). Second, patient tumors with high VEGFR2 staining had fewer CD8^+^ T cell and NK cell infiltrates (Fig. 4K-L). xCell analysis (55) of RNA-seq data from primary prostate tumor patient samples from the TCGA further confirmed these observations, where tumors with higher VEGFR2 (*KDR*) expression had fewer total as well as naïve and effector CD8^+^ T cell transcripts (Supplementary Fig. S4E). Finally, co-IF staining of prostate patient samples revealed a substantial percentage of CD8^+^ T cells that were VEGFR2^+^ in MYC^Hi^ tumors (Fig. 4M-N). Collectively, these data demonstrate that VEGF-VEGFR2 signaling orchestrated by *MYC* induction and *p53* inactivation in prostate cancers of mice and humans can directly suppress CD8^+^ T cytotoxicity and effector functions.

### VEGF signaling blockade reactivates anti-tumor CD8^+^ T cell immunity in *MP*-driven prostate cancer

As VEGFR2 blockade could enhance CD8^+^ T cell effector functions *in vitro*, we next asked whether VEGF signaling inhibition could remodel the immune suppressive landscape of *MP* tumors and restore anti-tumor T cell immunity *in vivo*. FVB mice were transplanted orthotopically with *Myc-p53KO* tumor cells, and upon tumor development, randomized into treatment groups where they received a VEGFR2 blocking antibody (DC101) or vehicle control. Following two-week treatment, prostate tumors were harvested, dissociated into single cell suspensions, and immunophenotyped by flow cytometry. α-VEGFR2 treatment led to a significant increase in both total CD3^+^ T cells, as well as cytotoxic CD8^+^ T cells (Fig. 5A). CD8^+^ T cells in the TME also had significantly higher levels of the activation markers IFNγ and GZMB following α-VEGFR2 treatment (Fig. 5B-C). Though NK cell numbers did not change upon VEGFR2 blockade, there was a trend toward increased effector functions as assessed by IFNγ and TNF⍺ expression (Fig. 5D). Moreover, suppressive FOXP3^+^ T_regs_ that inhibit CD8^+^ T cell and NK cell activity and accumulate preferentially in the *MP* genetic subtype of prostate cancer were also reduced by VEGFR2 blockade (Fig. 5E). Myeloid subsets, including macrophages, myeloid-derived suppressor cells (MDSCs), and dendritic cells (DCs), remained unaltered following treatment (Supplementary Fig. 5A).

**Figure 5.**
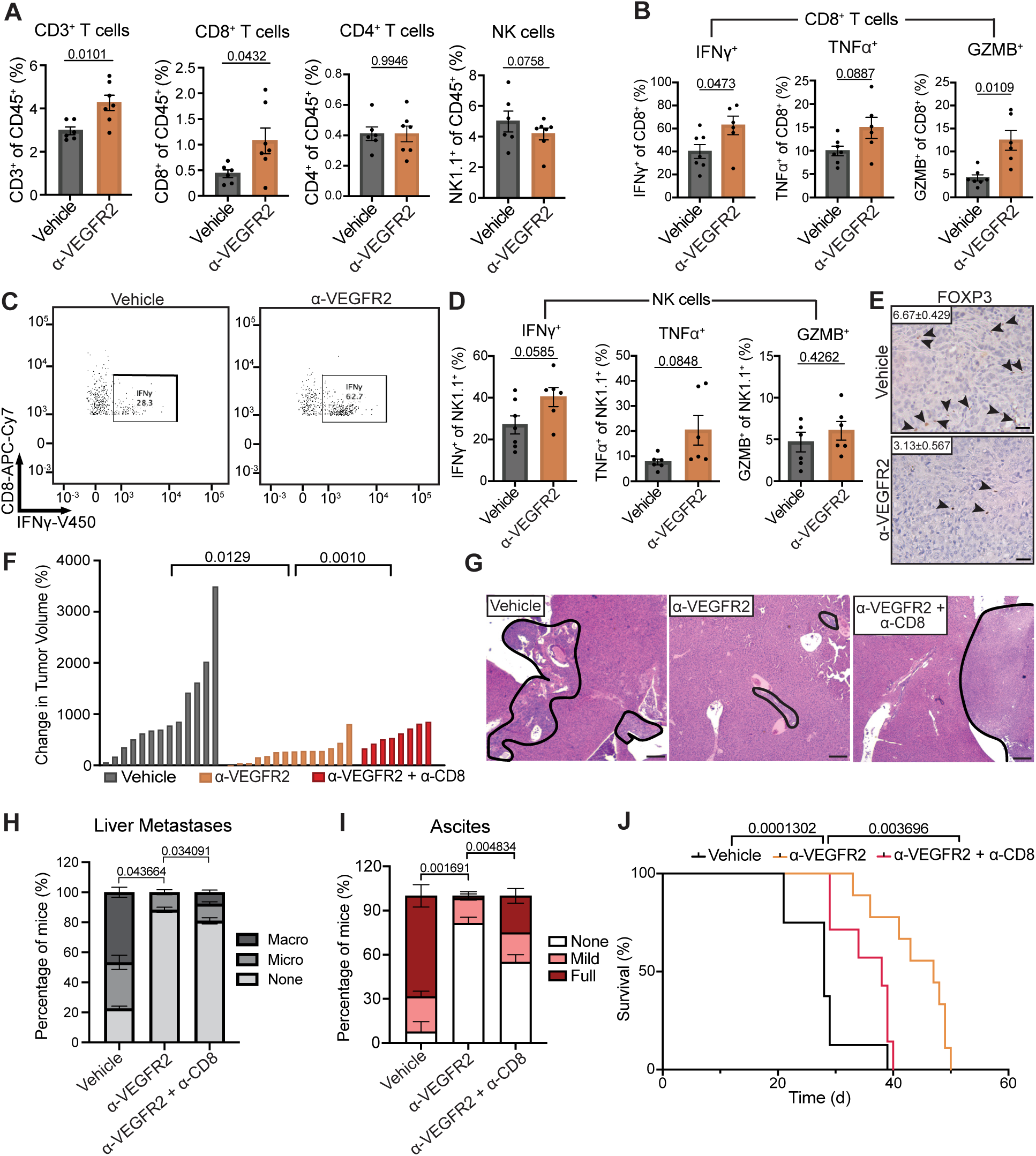
VEGFR2 neutralization can restore anti-tumor CD8^+^ T cell immunity in *MYC* and *p53* altered prostate cancer. **A,** Flow cytometry analysis of CD3^+^, CD8^+^, and CD4^+^ T cell, and NK1.1^+^ NK cell numbers in *Myc-p53*KO transplant prostate tumors from mice treated with vehicle or a VEGFR2 blocking antibody (DC101; 400μg) for 2 weeks (n = 6-7 mice per group). **B,** Flow cytometry analysis of expression of IFNγ, GZMB, and TNFα in CD8^+^ T cells in *Myc-p53*KO transplant prostate tumors from mice treated as in **A** (n = 6-7 mice per group). **C,** Representative flow cytometry plots of IFNγ expression in CD8^+^ T cells in *Myc-p53*KO transplant prostate tumors from mice treated as in **A**. **D,** Flow cytometry analysis of expression of IFNγ, GZMB, and TNFα in NK1.1^+^ NK cells in *Myc-p53*KO transplant prostate tumors from mice treated as in **A** (n = 6-7 mice per group). **E,** Representative IHC staining of *Myc-p53*KO transplant prostate tumors treated as in **A** and harvested at endpoint. Arrowheads indicate positive staining for immune cells. Quantification of FOXP3^+^ T_regs_ per field is shown inset (n = 5 mice per group). Scale bars, 50μm. **F,** Waterfall plot of response of *Myc-p53*KO transplant tumors to 2-week treatment with vehicle, α-VEGFR2 (DC101; 400μg), and/or a CD8 neutralizing antibody (2.43; 200μg) (n = 7-13 mice per group). **G,** Representative hematoxylin and eosin (H&E) staining of liver metastases (outlined in black) from *Myc-p53*KO prostate tumor-bearing mice treated as in **F**. Scale bars, 100μm. **H,** Quantification of the percentage *Myc-p53*KO prostate tumor-bearing mice with micro- or macrometastases in the liver at endpoint following treatment as in **F** (n = 7-9 per group). **I,** Quantification of percentage *Myc-p53*KO prostate tumor-bearing mice with mild or full ascites at endpoint following treatment as in **F** (n = 7-9 per group). **J,** Kaplan-Meier survival curve of *Myc-p53*KO prostate tumor-bearing mice treated as in **F** (n = 7-9 per group). Data represent mean ± SEM. P-values were calculated by two-tailed, unpaired Student’s t-test (**A,B,D,F,H,I**) and log-rank (**J**).

To investigate whether this remodeling of the tumor-immune landscape following VEGFR2 blockade was specific to *MP* tumors, we also performed the same experiment using transplanted *Myc-CaP* parental cells that have the p53 locus intact. We did not observe the same increase in T cell numbers and activation of CD8^+^ T cell responses in *Myc-CaP*-derived tumors that occurred in *Myc-p53*KO tumors following VEGFR2 antibody administration (Supplementary Fig. S5B-D). Moreover, there was no change in blood vessel density, lumen structure, or expression of endothelial activation markers such as ICAM-1 and VCAM-1 that are important for T cell extravasation after VEGFR2 antibody treatment of *Myc-p53*KO tumor-bearing mice, suggesting that the effects of VEGFR2 inhibition on immunity were likely independent of vascular remodeling (Supplementary Fig. 5E-H). Together these findings suggest that VEGFR2 blockade can directly activate CD8^+^ T cell responses in prostate tumors with compound *MYC* and *p53* alterations *in vivo*.

We further assessed the short- and long-term impact of VEGFR2 blockade on tumor growth, metastasis, and animal survival. *Myc-p53*KO prostate tumors from α-VEGFR2-treated mice had significantly reduced growth after two-week treatment compared to those from control vehicle-treated mice (Fig. 5F; Supplementary Fig. 5I). Moreover, α-VEGFR2 treatment resulted in a reduction in visceral metastases to liver, as well as the incidence of ascites that is a common result of metastatic seeding of tumor cells to peritoneum and other organs at endpoint (Fig. 5G-I; Supplementary Fig. 5J). This reduction in primary tumor as well as metastatic burden resulted in significantly enhanced overall survival of prostate tumor-bearing mice treated with VEGFR2 blocking antibodies (Fig. 5J). Finally, to determine if CD8^+^ T cells activated following treatment functionally contributed to the anti-tumor effects of VEGFR2 blockade, we also administered a CD8 depleting antibody (2.43) to some animals. CD8^+^ T cell ablation significantly reduced the survival benefits of α-VEGFR2 blockade and led to an increased tumor and metastatic burden following treatment (Fig. 5F-J). Thus, VEGF signaling blockade can produce a more “inflamed” TME in *MP*-driven CRPC that culminates in CD8^+^ T cell-mediated tumor control and increased survival outcomes.

### VEGFR2 blockade improves anti-PDL-1 ICB efficacy in preclinical CRPC models

The immune suppressive prostate TME devoid of cytotoxic lymphocytes is thought to contribute to *de novo* resistance to anti-PD1/PD-L1 and anti-CTLA-4 ICB that has been curative in other malignancies (9–12,56). Indeed, we found that anti-PD-1 (RMP1-14) regimens had no impact on tumor growth or survival in preclinical *MP*-driven prostate cancer models (Supplementary Fig.6A-D). Interestingly, we observed induction of PD-L1 expression in *Myc-p53*KO prostate tumors following VEGFR2 blockade (Fig. 6A), likely the downstream result of IFNγ production by activated T cells. This increase in activated T cells coupled with induction of PD-L1 on tumor cells following α-VEGFR2 administration provided strong rationale for combining VEGFR2 blocking antibodies with anti-PD-L1 ICB.

**Figure 6.**
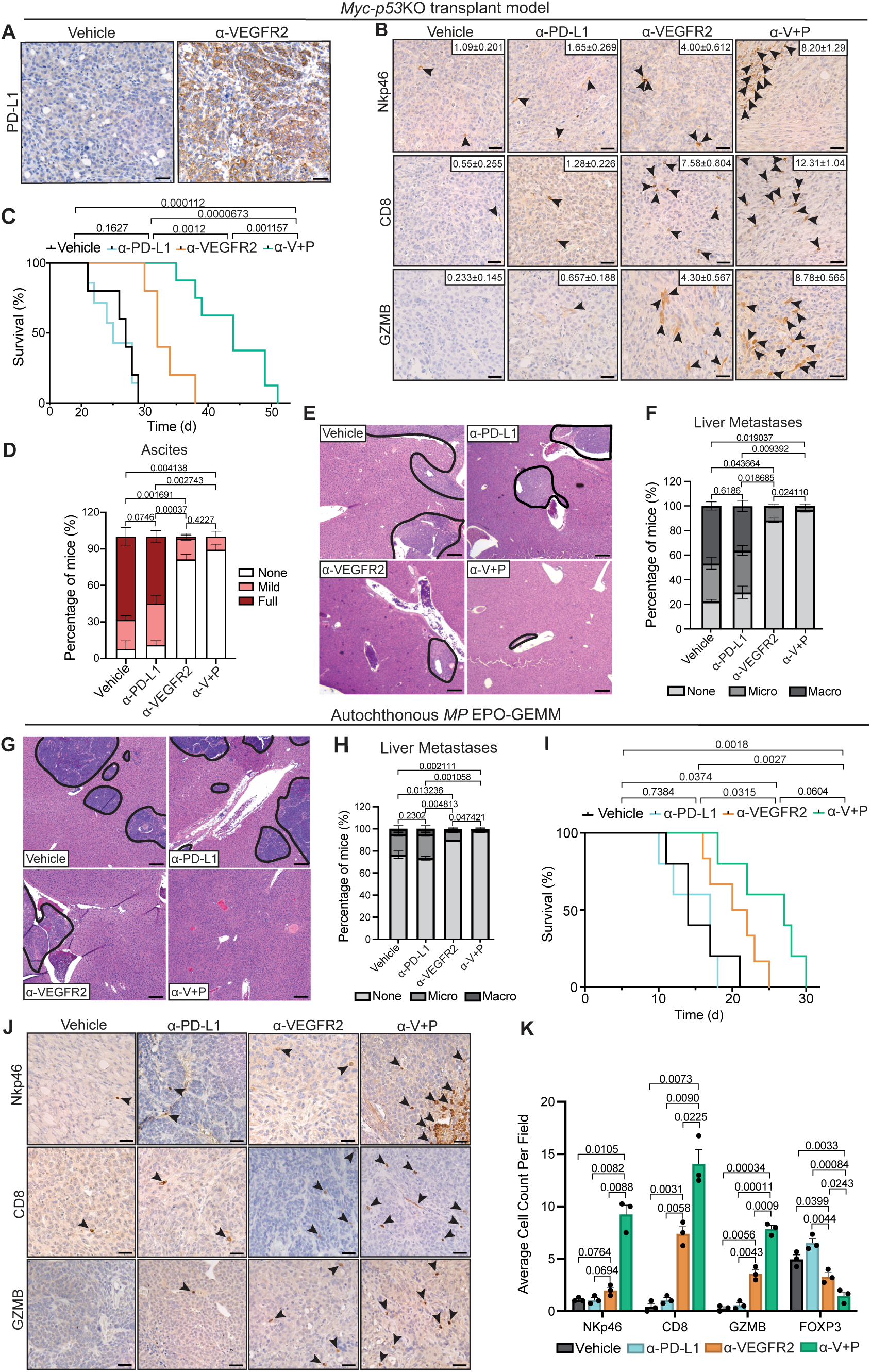
Combined VEGFR2 and anti-PD-L1 immune checkpoint blockade produces anti-tumor efficacy in preclinical prostate cancer models. **A,** Representative IHC staining for PD-L1 in *Myc-p53*KO transplant prostate tumors from FVB mice treated with vehicle or a VEGFR2 blocking antibody (DC101; 400μg) for 2 weeks. Scale bars, 50μm. **B,** Representative IHC staining for immune markers in *Myc-p53*KO transplant prostate tumors harvested at endpoint from mice treated with vehicle, VEGFR2 (V) (DC101; 400μg), and/or PD-L1 (P) (10F.9G2; 200μg) blocking antibodies. Arrowheads indicate positive staining for immune cells. Quantification of NKp46^+^ NK cells, CD8^+^ T cells, and GZMB^+^ cytotoxic lymphocytes per field is shown inset (n = 5 mice per group). Scale bars, 50μm. **C,** Kaplan-Meier survival curve of *Myc-p53*KO prostate tumor-bearing mice treated in **B** (n = 6-9 mice per group). **D,** Quantification of percentage of *Myc-p53*KO prostate tumor-bearing mice with mild or full ascites at endpoint following treatment as in **B** (n = 6-9 mice per group). **E,** Representative H&E staining of liver metastases (outlined in black) in *Myc-p53*KO prostate tumor-bearing mice treated as in **B**. Scale bars, 100μm. **F,** Quantification of percentage of *Myc-p53*KO prostate tumor-bearing mice with micro- or macrometastases in the liver at endpoint following treatment as in **B** (n = 6-9 mice per group). **G,** Representative H&E staining of liver metastases (outlined in black) in autochthonous prostate tumor-bearing *MP* EPO-GEMMs treated as in **B**. Scale bars, 100μm. **H,** Quantification of percentage of autochthonous prostate tumor-bearing *MP* EPO-GEMM animals with micro- or macrometastases in the liver at endpoint following treatment as in **B** (n = 4-6 mice per group). **I,** Kaplan-Meier survival curve of autochthonous prostate tumor-bearing *MP* EPO-GEMM animals treated as in **B** (n = 4-6 mice per group). **J,** Representative IHC staining for immune markers in autochthonous prostate tumors harvested at endpoint from *MP* EPO-GEMM mice treated as in **B**. Arrowheads indicate positive staining for immune cells. Scale bars, 50μm. **K,** Quantification of NKp46^+^ NK cells, CD8^+^ T cells, GZMB^+^ cytotoxic lymphocytes, and FOXP3^+^ T_regs_ per field from IHC staining in **J** (n = 3 mice per group). Data represent mean ± SEM. P-values were calculated by two-tailed, unpaired Student’s t-test (**D, F, H, K**) and log-rank (**C, I**).

We first treated FVB mice harboring transplanted *Myc-p53*KO prostate tumors with vehicle or antibodies targeting VEGFR2 or PD-L1 (10F.9G2) alone or in combination to assess the impact on tumor and immune responses by IHC. Consistent with its lack of efficacy as a monotherapy in patients, PD-L1 ICB had no impact on NK and CD8^+^ T cell frequencies and cytotoxicity, as well as on numbers of suppressive FOXP3^+^ T_regs_ in the prostate TME (Fig. 6B; Supplementary Fig. S6E). In contrast, combining PD-L1 with VEGFR2 blockade led to a significant increase in NK and CD8^+^ T cell accumulation and GZMB expression and reduction in T_regs_ in the prostate TME compared to not only PD-L1 ICB monotherapy but even to anti-VEGFR2 single agent treatment (Fig. 6B; Supplementary Fig. S6E). Animals were subsequently treated continuously with single or dual antibody treatment to assess the long-term effects on metastatic progression and survival. Whereas single arm α-PD-L1 dosing resulted in comparable animal survival to control vehicle treatment, combined VEGFR2 and PD-L1 blockade achieved robust and significant increases in overall survival compared to either antibody regimen alone (Fig. 6C). This survival advantage following dual VEGFR2/PD-L1 blockade also corresponded with a reduction in the presentation of ascites and metastases to the liver (Fig. 6D-F).

To further validate the preclinical efficacy and immune remodeling capacity of VEGFR2 and PD-L1 antibody treatment, we evaluated these regimens in autochthonous *MP* EPO-GEMM models. Treatment of prostate tumor-bearing *MP* EPO-GEMMs with the combination of VEGFR2 and PD-L1 blockade resulted in reduced primary tumor volumes and increased areas of tumor necrosis after two weeks of treatment, as well as diminished metastatic spread to the liver, compared to either single treatment alone (Fig. 6G-H, Supplementary Fig. 6F-G). These effects on primary and metastatic tumors culminated in greater overall survival for *MP* EPO-GEMMs treated with combined VEGFR2 and PD-L1 blockade compared to those receiving single agent treatment in these aggressive models (Fig. 6I). Importantly, we found that VEGFR2 blockade alone or in combination with PD-L1 ICB had similar effects on remodeling the immune suppressive prostate TME in autochthonous *MP* EPO-GEMM models as found in transplanted *Myc-p53*KO models. We observed an increased accumulation of NK cells and CD8^+^ T cells, induction of cytotoxic GZMB expression, and reduction in suppressive T_reg_ populations within the TME of *MP* EPO-GEMM tumors treated with α-VEGFR2 alone that was significantly enhanced following dual VEGFR2 and PD-L1 blockade (Fig 6J-K; Supplementary Fig. 6H). Collectively, our results demonstrate that VEGFR2 blockade can potentiate the effects of PD-L1 blockade to enhance anti-tumor immunity, extend overall survival, and even block metastatic progression in multiple aggressive and late-stage preclinical models of CRPC.

## DISCUSSION

For prostate cancer patients that relapse on hormone therapy and develop CRPC, there are still no durable treatment options. Immune checkpoint blockade (ICB) regimens that can produce curative responses in treatment-refractory melanoma and lung cancer have been generally ineffective in prostate malignancies, owing to their immune suppressed or “cold” tumor microenvironment (TME) that is devoid of cytotoxic T cells (9–12,56). Here we used an innovative *in vivo* electroporation approach to engineer genetic alterations that commonly arise in human CRPC, including amplification of *MYC* and deletion or mutation of tumor suppressor genes (TSGs) *P53*, *PTEN*, and *RB1*, in mouse models of prostate cancer. While *MYC* overexpression alone led to some suppression of cytotoxic T and NK lymphocytes, the most potent immune suppression was observed in combination with *p53* alterations. *MYC* and *p53* (*MP*) co-alterations in human prostate cancers were also associated with lymphocyte suppression and significantly reduced overall survival outcomes. Mechanistically, *MYC* induction and *p53* deficiency cooperated to promote tumor intrinsic secretion of VEGF, which can bind to its cognate receptor VEGFR2 that is expressed substantially on infiltrating T cells to inhibit their function (Fig. 7A). Treatment of *MP* prostate tumor-bearing mice with VEGFR2 blocking antibodies resulted in CD8^+^ T cell-mediated tumor and metastasis control. VEGF-VEGFR2 signaling inhibition also led to robust PD-L1 upregulation in prostate tumors, and combined VEGFR2 and PD-L1 antibody treatment produced significant anti-tumor T cell responses and survival outcomes in multiple aggressive and late-stage preclinical prostate cancer models resistant to ICB alone (Fig. 7B). As such, our results unveil VEGF-VEGFR2 signaling as a novel tumor intrinsic mechanism and biomarker of immune suppression in prostate cancer and promising target to potentiate immunotherapy in an aggressive subtype of CRPC lacking in effective treatment options.

**Figure 7.**
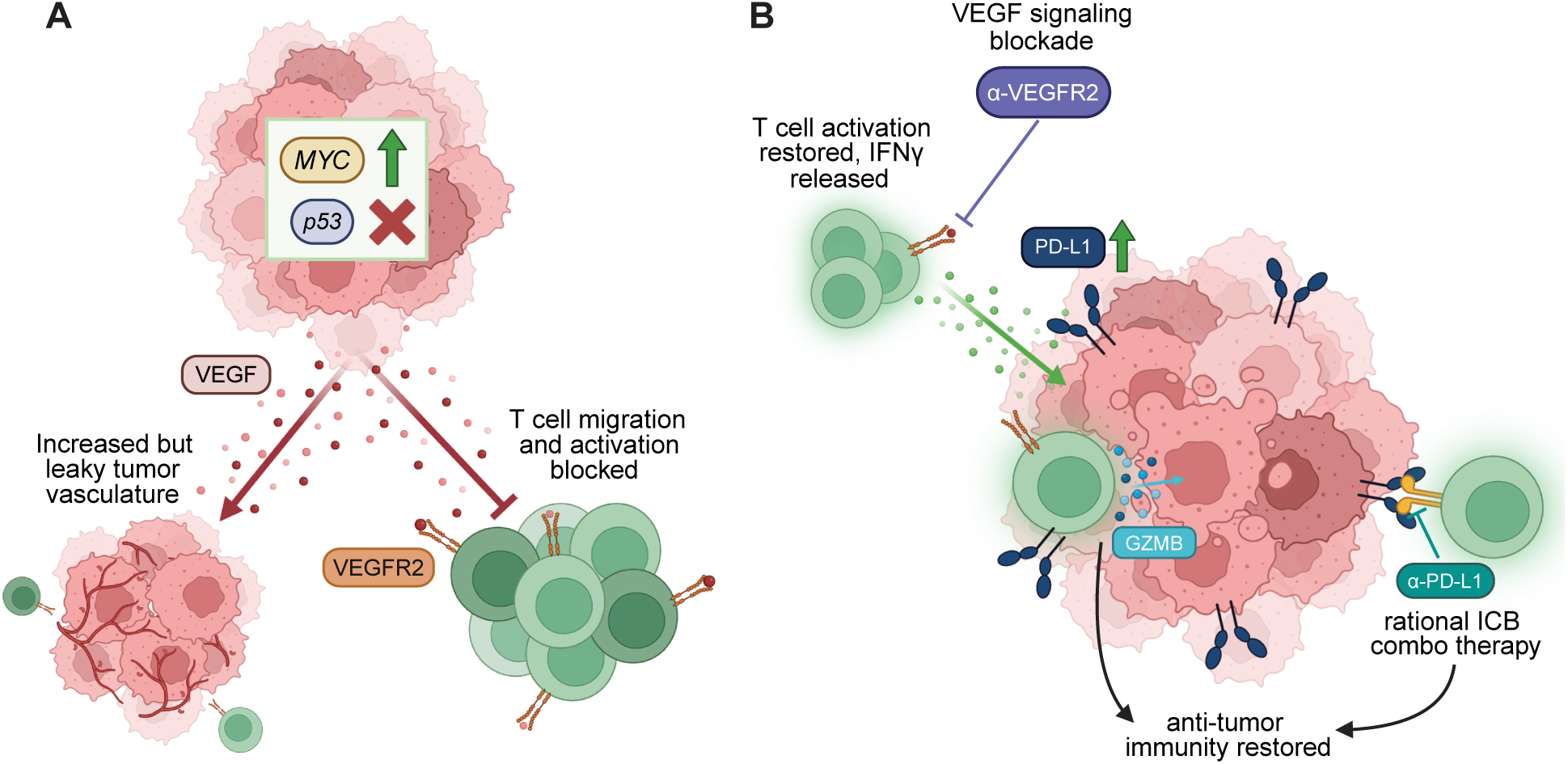
Tumor-derived VEGF signaling orchestrates an immune suppressive TME that can be overcome with combined VEGFR2 and PD-L1 blockade to restore T cell-mediated tumor control of *MYC* and *p53* co-altered prostate cancer. **A,** *MYC* overexpression and *p53* loss of function cooperate to promote secretion of VEGF, which leads not only to increased angiogenesis and a leaky vasculature, but also to direct inhibition of CD8^+^ T cells expressing VEGFR2 in the prostate TME. **B,** VEGFR2 inhibition enhances CD8^+^ T cell activity, and in combination with blockade of PD-L1 that is upregulated following treatment, produces potent anti-tumor efficacy in preclinical prostate cancer models. Figure created with Biorender.com.

MYC has long been described as a driver of oncogenesis, but more recently its role in generating an immunosuppressive TME has begun to surface (57). Our findings using an innovative *in vivo* genetic engineering approach further expand on this concept by demonstrating that while MYC induction can indeed suppress inflammatory signaling networks and NK and T cell responses in prostate cancer, this immune suppression is further enhanced in combination with *p53* disruption. Alterations in *MYC* and *p53* converge to induce expression of VEGF in cancer cells as well as VEGFR2 on T cells that directly inhibits CD8^+^ T cell function. In contrast to a lung adenocarcinoma study demonstrating that MYC, through induction of CCL9 expression, can recruit macrophages that secrete VEGF (34), we find that VEGF is directly produced by tumor cells rather than macrophages, which are not significantly changed in *MP* prostate tumors. As much previous work in prostate cancer has focused on the role of suppressive macrophages and MDSCs in orchestrating immune suppression (58,59), particularly in the context of *Pten* deletion (29,33), our results demonstrate an alternative tumor intrinsic mechanism of VEGF-mediated immune suppression that appears to be independent of myeloid cells. This suggests that there may exist distinct mechanisms of immune evasion in prostate cancer that are genetically driven and could be uniquely targetable for precision medicine. Still, the molecular mechanisms by which *MYC* induction and *p53* loss cooperate to drive VEGF expression warrant further investigation. Given that interferon signaling important for both MHC-I and PD-L1 expression is suppressed in *MYC* and *p53* co-altered tumors, along with previous literature indicating that VEGF can repress the Type I interferon receptor IFNAR1 and its downstream signaling (60–62), it will be of interest to explore the role of IFN signaling modulation in prostate cancer immune responses in future studies. Importantly, the EPO-GEMM platform can be leveraged to interrogate the role of other genes co-altered with *MYC* in prostate cancer (*BRCA1/2*, *APC, CHD1*) in mice with different host backgrounds (e.g. *NU/NU*, *Ifnar1^-^/^-^*) in order to further the dissect tumor-immune interactions in other genetic settings.

VEGF has a long-established role in regulating angiogenesis through activating VEGFR2 expressed on endothelial cells that fuels tumor invasion and metastasis (63). In addition, the leaky and poor vascular integrity mediated by chronic VEGF signaling can inhibit effective extravasation of T lymphocytes into tissues (64). Indeed, recent pan-cancer meta-analysis of angiogenic and immune signatures demonstrated that greater than 80% of prostate cancers are associated with an inversely related high angiogenic and low T cell activity score predictive of poor responses to ICB therapy (65). Vascular normalization through administration of low doses of antibodies targeting VEGF or VEGFR2 has been pursued as a strategy to increase immune cell infiltration and function, as well as potentiate immunotherapy responses in various cancer settings (66–68). Here, we find that though VEGF secretion is associated with an increased number of poorly formed blood vessels in tumors with *MYC* and *p53* co-alterations, the vasculature per se does not seem to directly contribute to immune suppression in our model as VEGFR2 blockade at the administered doses has no effect on blood vessel numbers or vascular phenotypes that could impact immune functions. In contrast, VEGF had a direct effect on the functions of CD8^+^ T cells expressing VEGFR2, leading to repression of effector cytokine (IFNγ) and GZMB secretion. Though there were no changes in the numbers of myeloid cells that can respond to or secrete VEGF, VEGFR2 blockade did significantly diminish the frequencies of regulatory T cells (T_regs_) that were enriched in *MP* tumors and whose suppression could also indirectly enhance effector CD8^+^ T cell function. Moreover, though not explored here, it is also possible that targeting of VEGFR2 expressed on tumors cells could also directly inhibit tumor growth. Future work using spatial imaging and transcriptomic approaches and ligand-receptor network analysis could provide deeper granularity into how VEGF-VEGFR2 signaling impacts different aspects of the immune suppressive TME of prostate cancer and whether our findings may apply to other cancer settings with high angiogenic activity.

VEGF signaling was first identified as a potential therapeutic target for solid cancer types over fifty years ago, and many attempts have been made since to block this pro-tumorigenic axis (69). In prostate cancer, despite VEGF expression being associated with disease progression and stage, neither VEGF neutralization with antibodies such as bevacizumab nor VEGFR2 inhibition through use of multi-receptor tyrosine kinase (RTK) inhibitors has led to a significant benefit in overall survival in combination with standard chemotherapy in phase III clinical trials in mCRPC patients (70,71). Still, some patients do initially respond to VEGF signaling blockade, indicating there may be a subset of patients where this treatment regimen could be effective (72,73). Our results indicate that the ∼8-9% of CRPC patients harboring co-alterations in *MYC* and *p53* may particularly benefit from clinically approved VEGF (e.g. bevacizumab) and VEGFR2 (e.g. Sunitinib) targeting therapies despite overall treatment failure in the broader population. Moreover, though this patient population presents with a severely immune suppressed TME and *de novo* resistance to PD-1/PD-L1 ICB, we find that VEGFR2 blockade induces robust PD-L1 expression and sensitivity to anti-PD-L1 ICB regimens in combination. Indeed, atezolizumab (anti-PD-L1) and bevacizumab (anti-VEGF) combinatorial therapy was recently FDA-approved for hepatocellular carcinoma (HCC), and preclinical studies demonstrate its effectiveness in a MYC-driven mouse model of HCC (74). Collectively, our results pave a clear translational path for the implementation of VEGFR2 and PD-L1 blocking antibodies clinically approved in other malignancies for the treatment of an aggressive CRPC subtype driven by *MYC* and *p53* alterations. More broadly, similar approaches could be taken to implement unique immunotherapy regimens based on the genetics of a tumor in prostate and other cancer types for “precision immunotherapy”.

## ACKNOWLEDGEMENTS

We thank J. Leibold for assistance with EPO-GEMM animal and cell line generation; J. Peura and J. Pitarresi for support with IHC/IF staining and imaging; S. Liu and K. Wagner for aid in initial project direction; M. Wang for providing murine prostate cancer cell lines and VEGF signaling expertise; J. Pitarresi and M. Kelliher for helpful comments and feedback on the manuscript; and members of the Ruscetti and Pitarresi labs for insightful feedback throughout the project. This work was supported by a Prostate Cancer Research Program (PCRP) Idea Development Award from the Department of Defense (DoD) office of the Congressionally Directed Medical Research Programs (CDMRP) (W81XWH-22-1-0505) to M.R., a National Center for Advancing Translational Sciences grant (UL1-TR001453) to K.S., and an R01 grant from the National Cancer Institute (NCI) (CA276863) to A.M.M.

## AUTHOR CONTRIBUTIONS

**Conceptualization:** K.C. Murphy, M. Ruscetti

**Data curation:** K.C. Murphy, J. Li, L.J. Zhu, Y.J. Ho

**Formal Analysis:** K.C. Murphy, J. Li, L.J. Zhu, Y.J. Ho

**Funding acquisition:** K. Simin, A.M. Mercurio, M. Ruscetti

**Investigation:** K.C. Murphy, K.D. DeMarco, L. Zhou, Y. Lopez, J. Li, S. Bai

**Methodology:** K.C. Murphy, K.D. DeMarco, L. Zhou, J. Li, Y.J. Ho

**Project administration:** K. Simin, L.J. Zhu, M. Ruscetti

**Resources:** K. Simin, L.J. Zhu, M. Ruscetti

**Software:** J. Li, L.J. Zhu, Y.J. Ho

**Supervision:** A.M. Mercurio, M. Ruscetti

**Validation:** K.C. Murphy, S. Bai

**Visualization:** K.C. Murphy, J. Li

**Writing – original draft:** K.C. Murphy, M. Ruscetti

**Writing – review & editing:** K.C. Murphy, A.M. Mercurio, M. Ruscetti

## METHODS

### Animal studies

All mouse experiments in this study were approved by the University of Massachusetts Chan Medical School Internal Animal Care and Use Committee (IACUC). Mice were maintained under specific pathogen-free conditions, and food and water were provided ad libitum. C57BL/6 and FVB male mice for transplantation models were purchased from Charles River Laboratories and Jackson Laboratory, respectively.

### Electroporation based non-germline genetically engineered mouse models (EPO-GEMMs)

The electroporation procedure was performed as previously described (38). Briefly, 8- to 12-week old WT C57BL/6 male mice were anesthetized with 2-3% isoflurane and a small incision made in the peritoneal cavity near the pelvic region. After locating one of the seminal vesicles and attached anterior lobe, 30 μL of plasmid mix (see specifications below) was injected into an anterior lobe of the prostate using a 27.5 gauge syringe. Tweezer electrodes were then placed around the injection bubble and two pulses of electrical current (60V) given for 35-millisecond lengths at 500-millisecond intervals were then applied using an *in vivo* electroporator (Nepa Gene NEPA21 Type II Electroporator). After electroporation, the peritoneal cavity was rinsed with 0.5 mL of prewarmed saline. The abdominal wall was then sutured with an absorbable Vicryl suture (Ethicon), and the skin was closed with wound clips (CellPoint Scientific Inc.) Mice were monitored for tumor development by palpation and ultrasound imaging. At study endpoint, prostate tumors were harvested and tissue divided for 10% formalin fixation for immunohistochemistry (IHC) or immunofluorescence (IF) analysis, or single cell suspensions for flow cytometry analysis.

To generate *MYC*; *p53^-/-^* (*MP*) EPO-GEMM tumors, 5μg of a pT3-MYC transposon vector, 1μg of Sleeping Beauty transposase (SB13), and 20μg of a pX330 CRISPR/Cas9 vector with an sgRNA targeting the *p53* locus (sequence: ACCCTGTCACCGAGACCCC) were injected into the anterior lobe of the prostate. To generate *MYC*; *Pten^-/-^*(*MPten*) EPO-GEMM tumors, 5μg of a pT3-MYC transposon vector, 1μg of SB13, and 20μg of a pX330 CRISPR/Cas9 vector with an sgRNA targeting the *Pten* locus (sequence: GTTTGTGGTCTGCCAGCTAA) were injected into the anterior lobe of the prostate. To generate *PtPRb* (*Pten^-/-^;p53^-/-^;Rb1^-/-^*) EPO-GEMM tumors, 20μg each of two pX330 CRISPR/Cas9 vectors, one harboring a sgRNA targeting the *p53* locus and another harboring tandem sgRNA sequences targeting *Pten* and *Rb1* (sequence: TGCGCGGGGTCGTCCTCCCG) were injected. The SB13 and pT3-EF1α transposon vector were a gift from Dr. Xin Chen at UCSF and pX330 vector a gift from Feng Zhang at the Broad Institute (Addgene #42230). Genome editing in resulting EPO-GEMM tumors was confirmed by Sanger sequencing.

### Cell lines

*MP*, *MPten,* and *PtPRb* murine prostate cancer cell lines were generated from EPO-GEMM tumors with these genotypes. EPO-GEMM prostate tumors were minced, digested in DMEM containing 3 mg/mL Dispase II (Gibco) and 1 mg/mL Collagenase IV (C5138;Sigma) for 1 hour at 37°C, and then plated on 10-cm culture dishes coated with 100 μg/mL collagen (PureCol; 5005; Advanced Biomatrix). Cells that attached to the plate were passaged at least three times to remove non-tumor cell contaminants. Sanger sequencing was performed to confirm that EPO-GEMM cell lines maintained the same genetic alterations as their respective EPO-GEMM tumors. *Myc-CaP* cells were obtained from A.M. Mercurio. All cell lines were cultured in a humidified incubator at 37°C with 5% CO_2_ and grown in DMEM supplemented with 10% FBS and 100 IU/ml penicillin/streptomycin (P/S). Cells used for *in vivo* transplantation experiments tested negative for mycoplasma.

### Clonogenic assays

Bicalutamide was purchased from Selleck Chemicals (S1190), dissolved in dimethyl sulfoxide (DMSO) to yield 10 mM stock solutions, and stored at −80 °C. EPO-GEMM-derived cell lines were treated with varying concentrations of bicalutamide (or DMSO as a vehicle control) for 7 days, with growth media with or without drugs changed every 3 days. The remaining cells were fixed with methanol (1%) and formaldehyde (1%), stained with 0.5% Crystal Violet and photographed using a digital scanner.

### CRISPR-mediated *p53 KO* in *Myc-CaP* cells

To knockout (KO) *p53*, *Myc-CaP* cells were transiently transfected using Lipofectamine™ 3000 Transfection Reagent (Thermo Fisher; L3000008) according to manufacturer’s protocol with 20 μg of a pX330 CRISPR/Cas9 construct containing a sgRNA targeting *p53* (sequence: ACCCTGTCACCGAGACCCC) or no sgRNA as a control. Cells with p53 deficiency were then selected by treatment with 10μM of the MDM2 inhibitor nutlin-3 (Selleck Chemicals; S1061) for 72 hours. Successful generation of *Myc-p53*KO cells was confirmed by loss of *p53* expression by RT-qPCR analysis.

### T cell co-culture assays

To isolate primary murine CD8^+^ T cells, spleens were harvested from male wildtype 8-10 week old C57BL/6 mice (for culturing with EPO-GEMM-derived cell lines) or FVB mice (for culturing with *Myc-CaP*-derived cell lines) and passed through a 70μm cell strainer. Cells were centrifuged at 1500 rpm x 5 minutes before red blood cells were then lysed with ACK lysis buffer (Quality Biological) for 5 minutes. Samples were centrifuged and then resuspended in FACS buffer (PBS supplemented with 2% FBS) before CD8^+^ T cells were isolated by negative selection using a CD8 T cell Isolation Kit according to the manufacturer’s protocol (Miltenyi Biotec; 130-104-075). Isolated CD8^+^ T cells were incubated in RPMI media supplemented with 10% FBS and stimulated for 1 hour with PMA (20 ng/ml, Sigma-Aldrich), Ionomycin (1 μg/ml, STEMCELL technologies), and monensin (2 μM, Biolegend) in a humidified incubator at 37°C with 5% CO_2_. CD8^+^ T cells were then added to a 96-well plate with 5×10^3^ prostate tumor cells in triplicate at an effector to target ratio of 10:1 with or without a VEGFR2 (DC101; 1μg/mL) blocking antibody. Some CD8^+^ T cells from C57BL/6 mice were directly exposed to 50ng/mL recombinant murine VEGF from R&D Systems (493-MV-005/CF) in the absence of tumor cell co-culture as a control condition. T cells were then incubated for 4 hours in a humidified incubator at 37°C with 5% CO_2_ before being trypsinized, resuspended in PBS supplemented with 2% FBS, and stained with cell surface antibodies against CD45 AF700 (30-F11; 1:320), CD3 BV650 (17A2; 1:300), CD8 FITC (53-6.7; 1:400), and VEGFR2 PE (AVAS12; 1:200) for 30 minutes at 4°C. To assess Granzyme B (GZMB), IFNγ, and TNFα levels in CD8^+^ T cells, intracellular staining was performed using the Foxp3/transcription factor staining buffer set (eBioscience), where cells were fixed, permeabilized, and then stained with GZMB APC (GB11, Biolegend; 1:100), IFNγ V450 (XMG1.2, TONBO Biosciences; 1:100), and TNFα PE-Cy7 (MP6-XT22, eBiosciences; 1:100) antibodies. GZMB, IFNγ, and TNFα positivity was evaluated by gating on CD3^+^CD8^+^ T cells on a FACSymphony A5 flow cytometer and analyzed using FlowJo (TreeStar). Flow cytometry gating strategies are shown in Supplementary Fig. S4C.

### Prostate orthotopic transplantation models

2.5×10^5^ *MP*, 5×10^5^ *MPten*, or 5×10^5^ *PtPRb* cells were resuspended in 15μl of Matrigel (Matrigel, BD) diluted 1:1 with cold DMEM/F12 media and transplanted into one anterior lobe of the prostate of 8-week-old C57BL/6 male mice. 1×10^6^ *Myc-CaP* or *Myc-p53*KO cells were resuspended in 15μl of Matrigel (Matrigel, BD) diluted 1:1 with cold DMEM/F12 media and transplanted into one anterior lobe of the prostate of 8-week-old FVB male mice. Following anesthetization using 2-3% isoflurane, an incision was made in the peritoneal cavity and the cell suspension was injected into an anterior lobe of the prostate using a Hamilton Syringe. The injection’s success was confirmed by the presence of a fluid bubble without any indications of leakage into the abdominal cavity. The abdominal wall was sutured with an absorbable Vicryl suture (Ethicon), and the skin was closed with wound clips (CellPoint Scientific Inc.). Mice were monitored for tumor development by ultrasound imaging and randomized into treatment groups upon tumor formation based on tumor volume. Following sacrifice, a portion of the prostate tumor tissue was preserved in 10% formalin for fixation, while another portion was used for flow cytometry analysis.

### *In vivo* blocking antibody administration

To assess the impact of VEGFR2 and/or PD-1/PD-L1 antibody blockade on tumor and immune responses and overall animal survival, mice harboring genetically-defined EPO-GEMM or transplanted prostate tumors were randomized based on tumor size into different treatment cohorts and received vehicle (PBS), αVEGFR2 (DC101; 400μg), αPD-L1 (10F.9G2; 200μg), αPD-1 (RMP1-14; 200μg), or combined VEGFR2 and PD-L1 blocking antibodies concurrently by intraperitoneal (i.p.) injection twice per week. To determine the impact of CD8^+^ T cell depletion on tumor progression and animal survival, mice were injected i.p. with an αCD8 (200 μg; 2.43) depleting antibody twice per week. Antibodies were purchased from BioXcell and diluted in PBS.

### Ultrasound imaging

High-contrast ultrasound imaging was performed on a Vevo 3100 System with a MS250 13- to 24-MHz scanhead (VisualSonics) to stage and quantify prostate tumor burden. Tumor volume was analyzed using Vevo 3100 software, version 5.50.

### Flow cytometry

For analysis of MHC-I expression in prostate cancer cell lines cultured *in vitro*, cells were trypsinized, resuspended in PBS supplemented with 2% FBS, and stained with an H-2Kb antibody (AF6-88.5.5.3, eBioscience; 1:200) for 30 minutes on ice. Flow cytometry was performed on a FACSymphony A5 cytometer, and data were analyzed using FlowJo (TreeStar).

To prepare single cell suspensions from *in vivo* tumor samples for flow cytometry analysis, tumors were minced with scissors into small pieces and placed in 5ml of collagenase buffer [1x HBSS w/ calcium and magnesium (GIBCO), 1 mg/ml Collagenase A (Roche) and 0.1 mg/ml DNaseI (DN25; Sigma)]. Samples were then transferred to C tubes and processed using program 37C_multi_A on a gentleMACS Octo dissociator with heaters (Miltenyi Biotec). Dissociated tissue was passed through a 70μm cell strainer and centrifuged at 1500 rpm x 5 minutes. Red blood cells were then lysed with ACK lysis buffer (Quality Biological) for 5 minutes and samples were centrifuged and then resuspended in FACS buffer (PBS supplemented with 2% FBS). Samples were incubated with the following antibodies for 30 minutes at 4°C: CD45 AF700 (30-F11; 1:320), NK1.1 BV605 (PK136; 1:200), CD3 BV650 (17A2; 1:300), CD8 PE-Cy7 (53-6.7; 1:400), CD4 PE-Cy5 (GK1.5; 1:200), F4/80 APC (BM8; 1:200), Gr-1 Pacific Blue (RB6-8C5; 1:200), CD11c FITC (N418; 1:200), MHC-II PE (M5/114.15.2; 1:200) (Biolegend) and CD11b BUV395 (M1/70; 1:1,280) (BD Biosciences). DAPI was used to distinguish live/dead cells. Flow cytometry was performed on BD LSR II and FACSymphony A5 cytometers. CD4^+^ and CD8^+^ CD3^+^ T cell, CD3^-^ NK1.1^+^ NK cell, CD11b^+^ F4/80^+^ macrophage, CD11c^+^CD11b^-^MHC-II^+^ and CD11c^+^CD11b^+^MHC-II^+^ dendritic cell, and CD11b^+^Gr-1^+^ MDSC numbers were analyzed using FlowJo (TreeStar).

To analyze Granzyme B (GZMB), IFNγ, and TNFα levels in NK and T cells, single cell suspensions from tumor tissue were resuspended in RPMI media supplemented with 10% FBS and 100 IU/ml P/S and incubated for 4 hours with PMA (20 ng/ml, Sigma-Aldrich), Ionomycin (1 μg/ml, STEMCELL technologies), and monensin (2 μM, Biolegend) in a humidified incubator at 37°C with 5% CO_2_. Cell surface staining was first performed with CD45 AF700 (30-F11; 1:320), NK1.1 BV605 (PK136; 1:200), CD3 BV650 (17A2; 1:300), CD8 APC-Cy7 (53-6.7; 1:200), and CD4 PE-Cy5 (GK1.5; 1:200) (Biolegend) antibodies. Intracellular staining was then performed using the Foxp3/transcription factor staining buffer set (eBioscience), where cells were fixed, permeabilized, and then stained with GZMB APC (GB11, Biolegend; 1:100), IFNγ V450 (XMG1.2, TONBO Biosciences; 1:100), and TNFα PE-Cy7 (MP6-XT22, eBiosciences; 1:100) antibodies. GZMB, IFNγ, and TNFα positivity was evaluated by gating on CD3^-^NK1.1^+^ NK cells and CD3^+^CD8^+^ T cells on a BD LSR II or FACSymphony A5 flow cytometer and analyzed using FlowJo (TreeStar) as described above.

### Cytokine array

Murine prostate cancer cells were cultured in a humidified incubator at 37°C with 5% CO_2_ and grown in fresh DMEM supplemented with 100 IU/ml penicillin/streptomycin (P/S). Conditioned media was then collected after 72 hours of culturing and cells trypsinized and counted using a Countess II cell counter (Invitrogen). Media samples were normalized based on cell number by diluting with culture media. 60µl aliquots were analyzed using a multiplex immunoassay (Mouse Cytokine/Chemokine 44-Plex) from Eve Technologies.

### RT-qPCR

Total RNA was isolated from mouse prostate cell lines or WT prostate tissue from 8-10 week old C57BL/6 mice using the RNeasy Mini Kit (Qiagen). Complementary DNA (cDNA) was synthesized using the TaqMan reverse transcription reagents (Applied Biosystems) according to the manufacturer’s instructions. Real-time qPCR was performed in triplicate using SYBR Green PCR Master Mix (Applied Biosystems) on the StepOnePlus Real-Time PCR system (Applied Biosystems). Gene expression values were calculated using the ΔΔCT method and normalized to *Gapdh* levels as an endogenous reference gene. Primer sequences are listed in Supplementary Table S1.

### Bulk RNA-seq analysis of EPO-GEMM prostate tumors

RNA-seq analysis was performed on bulk prostate tumors from *MP*, *MPten*, and *PtPRb* EPO-GEMM mice as previously described (38). Heatmaps were generated using pheatmap. Gene set enrichment analysis (GSEA) was performed using the GSEAPreranked tool against Hallmark gene sets.

### Immunohistochemistry (IHC)

Murine prostate tissues were fixed overnight in 10% formalin and paraffin embedded. Formalin-fixed, paraffin-embedded (FFPE) blocks were then cut into 5μm sections. Hematoxylin and eosin (H&E) and IHC staining were performed using standard protocols. Sections were de-paraffinized, rehydrated, and boiled in a pressure cooker for 15 minutes in 10mM citrate buffer (pH 6.0) or 10 mM Tris base, 1 mM EDTA, 0.05% Tween 20 buffer (pH 9.0) for antigen retrieval. Endogenous peroxidases were quenched by incubating the slides in 3% hydrogen peroxide for 15 minutes. The sections were then washed 2x with PBS and blocked for 1 hour in 5% bovine serum albumin (BSA) in PBS solution at room temperature. Tissues were incubated overnight at 4°C in primary antibodies at respective dilutions (see Supplementary Table S2). HRP-conjugated secondary antibodies (Vectastain ImmPRESS®: Rabbit, MP-7401-50; Mouse, MP-7402-15; Goat, MP-7405-15; Rat, MP-7444-15) were then applied for 30 minutes and visualized with DAB (Vector Laboratories; SK-4100). Images were obtained on an Aperio ScanScope (Leica Biosystems). For immune cell and blood vessel quantifications, 10-20 high power 20x fields per section were counted and averaged using ImageScope v.12.3.2.8013 software from Leica Biosystems.

### Immunofluorescence (IF)

Prostate tissue sections were prepared for IF staining using standard protocols as described for IHC. Primary antibodies were incubated overnight at 4°C (see Supplementary Table S2). Secondary Alexa Fluor 488 or 647 dye-conjugated antibodies (Thermo Fisher; 1:150) were then applied for 1 hour at room temperature. Slides were mounted with Prolong Gold Antifade mountant (Prolong Molecular Probes; P36934) after counterstaining with DAPI. Fluorescent images were obtained on a Zeiss Axio Observer 7 microscope and quantified using Fiji.

### Analysis of liver metastasis and ascites burden

The incidence of metastasis to the liver was determined at study endpoint by analysis of H&E-stained liver sections at 10x magnification on an Aperio ScanScope (Leica Biosystems) in a blinded manner by K.C. Murphy. Micrometastases were defined as <100 cells and macrometastases >100 cells. The presence of ascites in prostate tumor-bearing mice was assessed at study endpoint based on mild (<500µL) or full (>500µL) amounts of bloody liquid in the peritoneal cavity.

### Prostate cancer patient samples

15 human prostate cancer specimens, including 5 of Gleason Score 6, 5 of Gleason Score 7, 2 of Gleason Score 8, and 3 of Gleason Score 9, were obtained from the UMass Center of Clinical and Translational Sciences Biorepository and derived retrospectively from patients undergoing surgery at UMass Memorial Hospital consented under the IRB approved protocol no. H-4721. De-identified FFPE tumor specimens were cut into 5μm sections and IHC staining performed as above (see Supplementary Table S2 for primary antibody information). IHC staining score grading was carried out by clinical pathologist B. Shi in a blinded manner. MYC staining was scored as low (<50% positive staining in tumor area) or high (>50% positive staining in tumor area) and VEGFR2 staining was scored on a 0-2 scale corresponding to low (little to no positive staining), intermediate (staining in some but not all tumor areas), and high (staining in majority of tumor areas). CD8^+^ T cell and NKp46^+^ NK cell numbers were counted by K.C. Murphy in a blinded manner and averaged from 20 high power 20x fields using ImageJ software.

### Human clinical data analysis

CBioPortal.org was used to construct an OncoPrint plotting the frequency of alterations in *MYC, P53, PTEN,* and *RB1* in mCRPC patients and generate a Kaplan-Meier survival curve of mCRPC patients harboring *MYC, P53,* and/or *PTEN* alterations from a SU2C dataset (15).

To analyze established immune signatures in prostate cancer patients with specific genetic alterations, we obtained gene expression dataset from The Cancer Genome Atlas (TCGA) (42) and corresponding genomic mutation datasets (75,76) from cBioPortal.org. Samples were categorized into groups based on the genomic status of *MYC, TP53*, and *PTEN*. Boxplots and Wilcoxon rank sum test were employed to compare the expression of inflamed/NK/T cell signatures (43–45) between groups using the R packages ggplot2 and ggpubr.

For xCell analysis of immune-related transcripts in tumors stratified by high and low expression of *KDR* (gene encoding VEGFR2), we downloaded the prostate adenocarcinoma expression dataset from The Pan-Cancer Atlas at gdc.cancer.gov. Using the R package xCell (55), we conducted cell type enrichment analysis for 64 immune and stromal cell types. Based on these results, violin plots were generated and Wilcoxon rank sum tests were performed on two distinct groups of samples stratified by the mean expression of *KDR*, utilizing the R packages ggplot2 and ggpubr for graphic representation and statistical testing.

### Statistical analysis

Statistical analyses were performed as described in the figure legend for each experiment. The indicated sample size (*n*) represents biological replicates and measurements were taken from distinct samples. No statistical method was used to predetermine sample size. Scoring of IHC/IF staining in mouse and human tumor samples was performed in a blinded manner. For other experiments, data collection and analysis were not performed blind to the conditions of the experiments. All samples that met proper experimental conditions were included in the analysis. Statistical significance was determined by Student *t* test, Wilcoxon test, or log-rank test using Prism 10 Software (GraphPad Software) or R as indicated.

## Data Availability

Bulk RNA-seq data generated or mined in this study are deposited in the Gene Expression Omnibus (GEO) database under accession numbers GSE139340 and GSE271975 (access token: arefoicojpkhrsl). All other data supporting the findings of this study are available from the corresponding author upon reasonable request.

