## Supplementary Figure S1 for "MYC and p53 alterations cooperate through VEGF signaling to repress cytotoxic T cell and immunotherapy responses in prostate cancer"

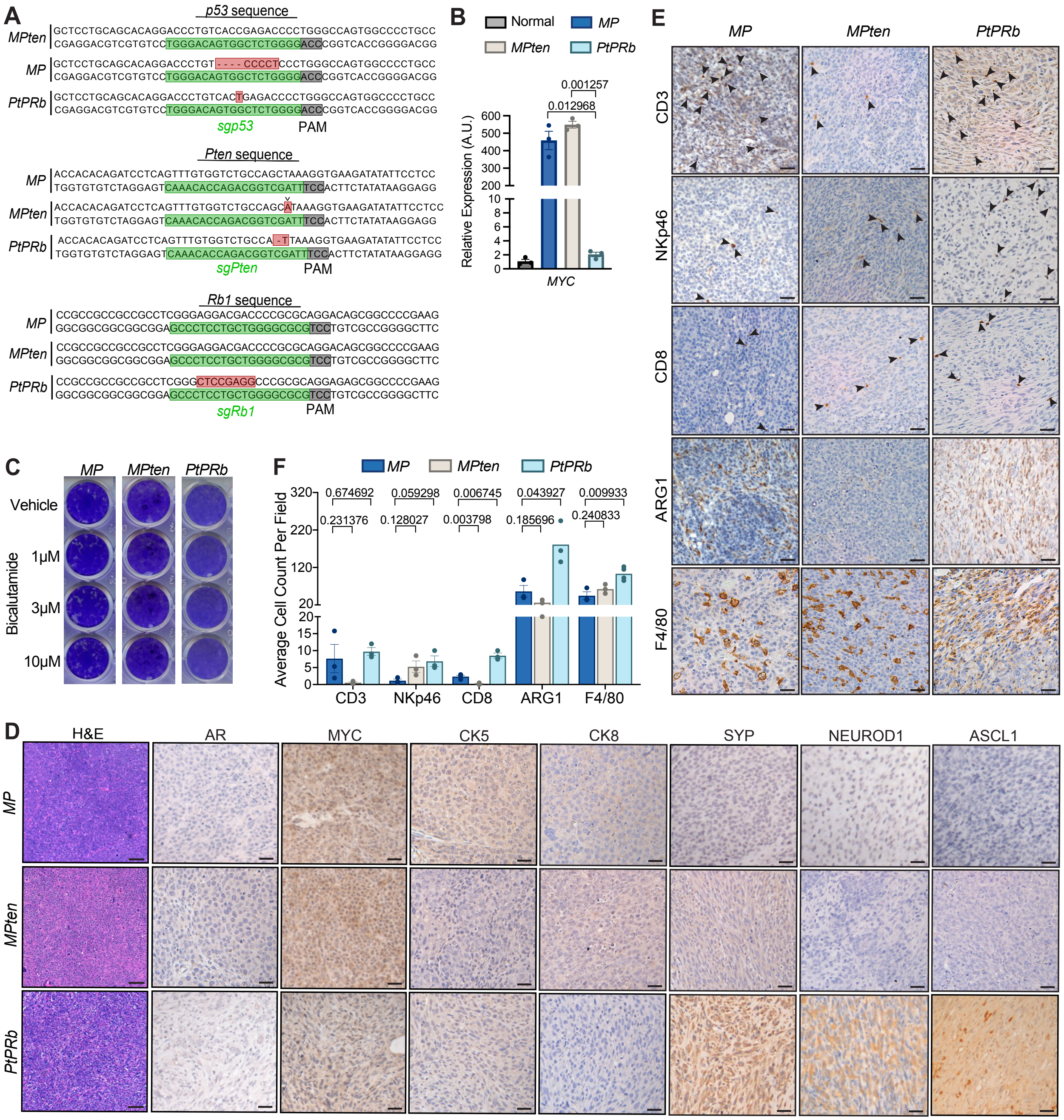


**Figure S1 (related to Figure 1). Genetic subtypes of murine AVPC display different immune milieus. A,** Sanger sequencing results showing editing of endogenous *P53*, *Pten*, and *Rb1* loci in *MP*, *MPten*, and *PtPRb* EPO-GEMM-derived prostate tumor cell lines. Red boxes indicate resulting indel from Cas9 editing. **B**, RT-qPCR analysis of human MYC expression in *MP*, *MPten*, and *PtPRb* EPO-GEMM-derived prostate tumor cell lines or control Wild-type (WT) prostate tissue from 8-10 week old C57BL/6 mice (n= 3 biological replicates). A.U., arbitrary units. **C,** Representative clonogenic assay images of *MP*, *MPten*, and *PtPRb* EPO-GEMM-derived prostate tumor cell lines treated with bicalutamide at indicated concentrations or a vehicle control (DMSO) for 7 days. **D,** Representative hematoxylin and eosin (H&E) and immunohistochemical (IHC) staining for MYC and prostate lineage markers in *MP*, *MPten*, and *PtPRb* EPO-GEMM prostate tumors harvested at endpoint. Scale bars, 50μm. **E,** Representative IHC staining of *MP*, *MPten*, and *PtPRb* EPO-GEMM-derived cell line transplant tumors harvested at endpoint. Arrowheads indicate positive staining for immune cells. Scale bars, 50μm. **F,** Quantification of $CD3^+$ T cells, $NKp46^+$ NK cells, $CD8^+$ T cells, $Arginase1^+$ (ARG1) suppressive myeloid cells, and $F4/80^+$ macrophages per field in **E** (n = 3-4 mice per group). Data represent mean ± SEM. P-values were calculated by two-tailed, unpaired Student's t-test.
