## Supplementary Figure S6 for "MYC and p53 alterations cooperate through VEGF signaling to repress cytotoxic T cell and immunotherapy responses in prostate cancer"

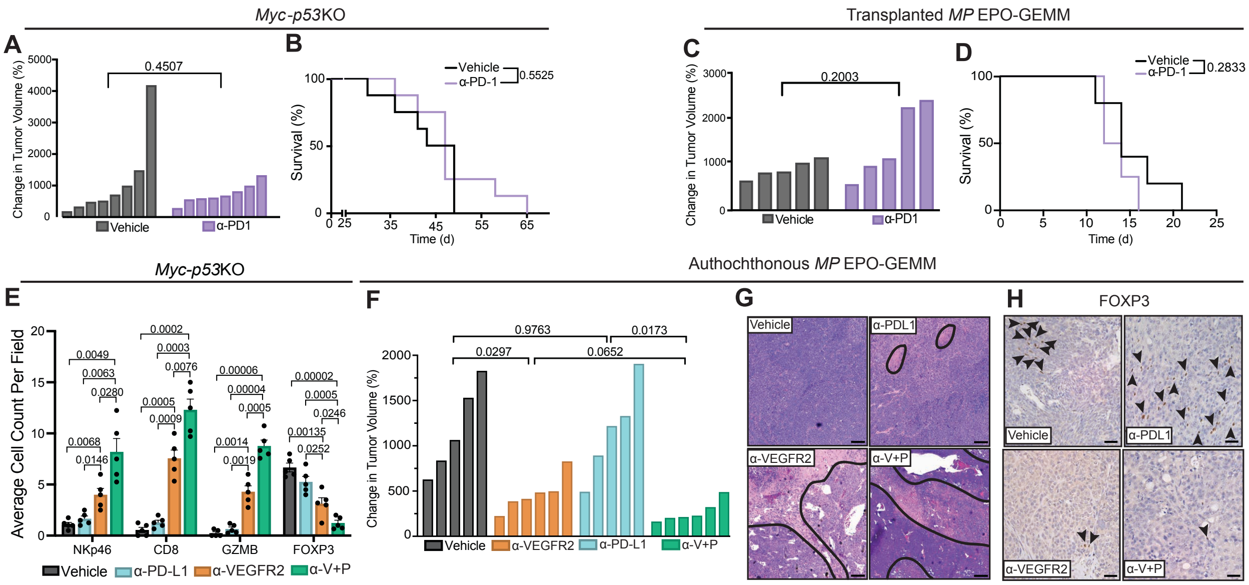


**Figure S6 (related to Figure 6). VEGFR2 targeting overcomes resistance to immune checkpoint blockade in *MYC* and p53 altered CRPC. A,** Waterfall plot of response of *Myc-p53*KO transplant prostate tumors to 2-week treatment with vehicle or α-PD-1 (RMP1-14; 200μg) (n = 8 mice per group). **B**, Kaplan-Meier survival curve of *Myc-p53*KO prostate tumor-bearing FVB mice treated as in **A** (n = 8 mice per group). **C,** Waterfall plot of response of transplanted *MP* EPO-GEMM prostate tumors to 2-week treatment as in **A** (n = 5 mice per group). **D,** Kaplan-Meier survival curve of C57BL/6 mice bearing transplanted *MP* EPO-GEMM prostate tumors treated as in **A** (n = 5 mice per group). **E,** Quantification of NKp46$^+$ NK cells, CD8$^+$ T cells, GZMB$^+$ cytotoxic lymphocytes, and FOXP3$^+$ $T_{regs}$ per field in *Myc-p53*KO transplant prostate tumors harvested at endpoint from mice treated with vehicle, VEGFR2 (V) (DC101; 400μg), and/or PD-L1 (P) (10F.9G2; 200μg) blocking antibodies from IHC staining in Fig. **6B** (n = 5 mice per group). **F,** Waterfall plot of response of autochthonous *MP* EPO-GEMM prostate tumors to 2-week treatment as in **E** (n = 5-6 mice per group). **G,** Representative H&E staining of autochthonous prostate tumors from *MP* EPO-GEMM mice treated as in **E**. Necrotic areas are outlined in black. **H,** Representative IHC staining in autochthonous *MP* EPO-GEMM prostate tumors harvested at endpoint from mice treated as in **E**. Arrowheads indicate positive staining for FOXP3$^{+}$ $T_{regs}$. Scale bars, 50μm. Data represent mean ± SEM. P-values were calculated by two-tailed, unpaired Student's t-test (**A, C, E, F**) and log-rank test (**B, D**).
