## Supplementary Figure S5 for "MYC and p53 alterations cooperate through VEGF signaling to repress cytotoxic T cell and immunotherapy responses in prostate cancer"

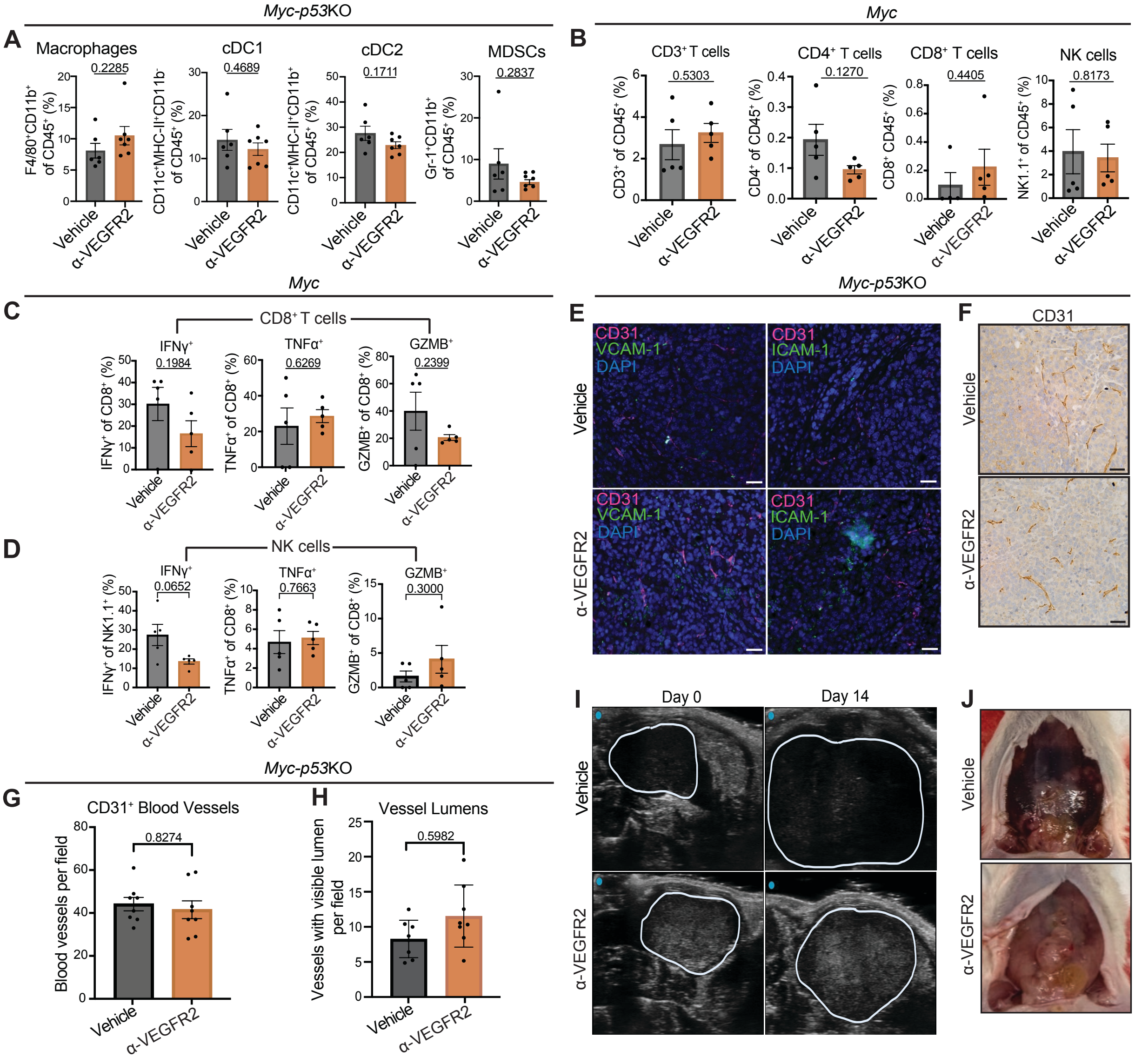


**Figure S5 (related to Figure 5). VEGFR2 blockade potentiates T cell-mediated prostate tumor and metastasis suppression specifically in *MP* tumors and independent of its effect on endothelial and myeloid cell phenotypes. A,** Flow cytometry analysis of F4/80$^+$ macrophage, CD11c$^+$MHC-II$^+$CD11b$^-$ cDC1 and CD11c$^+$MHC-II$^+$CD11b$^+$ cDC2 dendritic cell, and GR1$^+$CD11b$^+$ myeloid-derived suppressor cell (MDSC) numbers in *Myc-p53*KO transplant prostate tumors from mice treated with vehicle or a VEGFR2 blocking antibody (DC101; 400µg) for 2 weeks (n = 6-7 mice per group). **B,** Flow cytometry analysis of CD3$^+$, CD4$^+$, and CD8$^+$ T cell, and NK1.1$^+$ NK cell numbers in *Myc-CaP* (*Myc*) transplant prostate tumors from mice treated as in **A** (n = 5 mice per group). **C-D,** Flow cytometry analysis of expression of IFNγ, GZMB, and TNFα in CD8$^+$ T cells **(C)** and NK cells **(D)** in *Myc-CaP* (*Myc*) transplant prostate tumors from mice treated as in **A** (n = 5 mice per group). **E,** Representative co-IF staining of VCAM-1 or ICAM-1 expression in CD31$^+$ blood vessels in *Myc-p53*KO transplant prostate tumors treated as in **A**. Scale bars, 50μm. **F,** Representative IHC staining for CD31 in *Myc-p53*KO transplant prostate tumors treated as in **A**. Scale bars, 50μm. **G-H,** Quantification of CD31$^+$ blood vessel numbers (**G**) and vessels with visible lumens (**H**) in *Myc-p53*KO transplant prostate tumors from mice treated as in **A** (n = 8 mice per group). Scale bars, 100μm. **I,** Representative ultrasound images of *Myc-p53*KO transplant prostate tumors prior to (Day 0) and after 2 weeks of treatment as in **A**. Tumors are outlined in white. **J,** Representative images of ascites in *Myc-p53*KO transplant prostate tumor-bearing mice treated as in **A**. Data represent mean ± SEM. P-values were calculated by two-tailed, unpaired Student's t-test.
