## Supplementary Figure S4 for "MYC and p53 alterations cooperate through VEGF signaling to repress cytotoxic T cell and immunotherapy responses in prostate cancer"

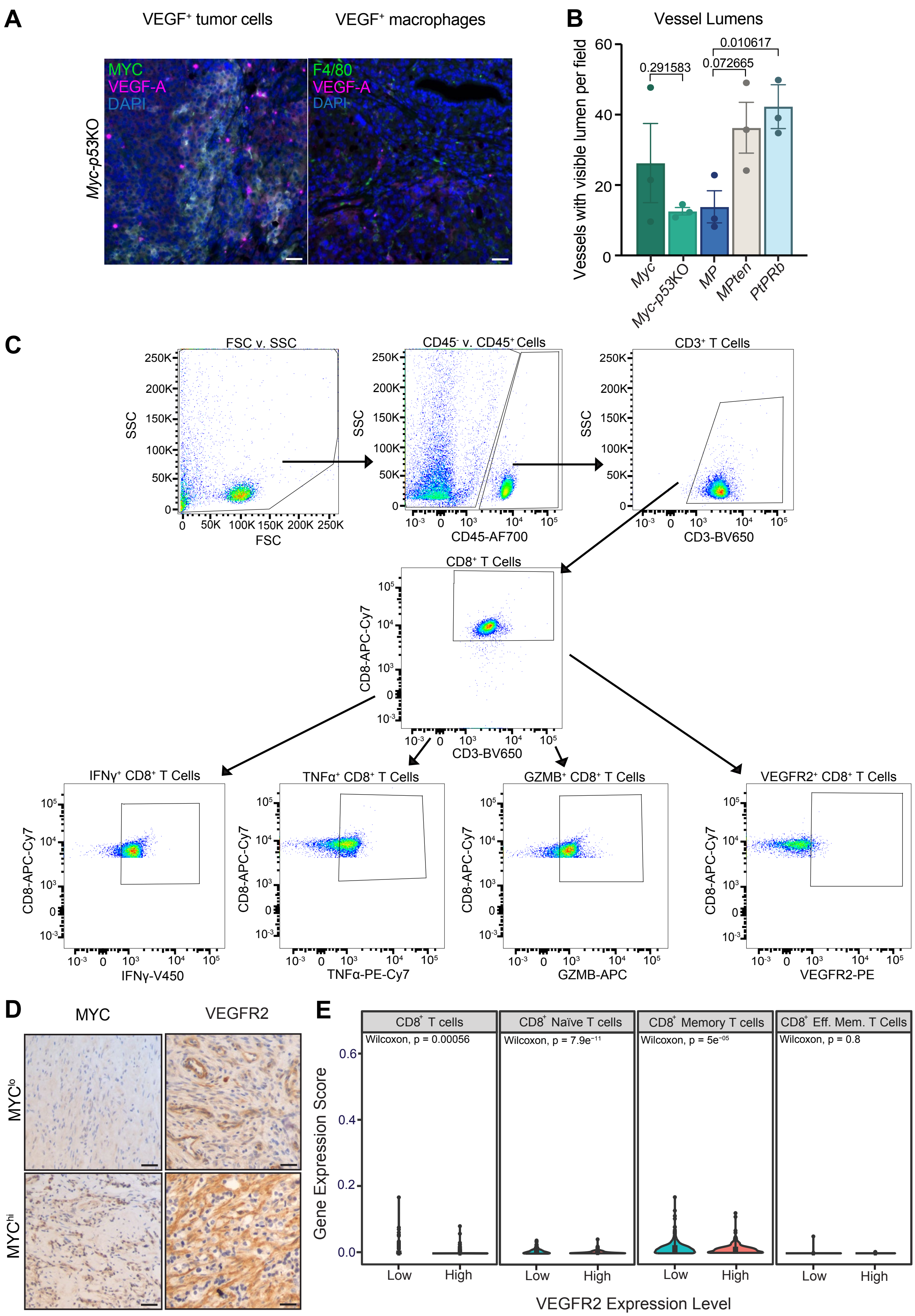

A
VEGF⁺ tumor cells
VEGF⁺ macrophages
Myc-p53KO
MYC
VEGF-A
DAPI
F4/80
VEGF-A
DAPI
B
Vessel Lumens
Vessels with visible lumen per field
0.291583
0.072665
0.010617
60
40
20
0
Myc
Myc-p53KO
MP
MPten
PtPRb
C
FSC v. SSC
SSC
FSC
CD45⁻ v. CD45⁺ Cells
CD45-AF700
CD3⁺ T Cells
CD3-BV650
CD8⁺ T Cells
CD8-APC-Cy7
CD3-BV650
IFNγ⁺ CD8⁺ T Cells
IFNγ-V450
TNFα⁺ CD8⁺ T Cells
TNFα-PE-Cy7
GZMB⁺ CD8⁺ T Cells
GZMB-APC
VEGFR2⁺ CD8⁺ T Cells
VEGFR2-PE
D
MYC
VEGFR2
MYClo
MYChi
E
CD8⁺ T cells
Wilcoxon, p = 0.00056
CD8⁺ Naïve T cells
Wilcoxon, p = 7.9e⁻¹¹
CD8⁺ Memory T cells
Wilcoxon, p = 5e⁻⁰⁵
CD8⁺ Eff. Mem. T Cells
Wilcoxon, p = 0.8
Gene Expression Score
0.6
0.4
0.2
0.0
Low
High
VEGFR2 Expression Level

**Figure S4 (related to Figure 4). *MYC*-driven prostate tumors have higher tumor intrinsic VEGF levels and VEGFR2 expression in the TME that are associated with T cell suppression.** **A,** Representative co-immunofluorescence (co-IF) staining of VEGF expression in $MYC^+$ tumor cells (left) or $F4/80^+$ macrophages (right) in *Myc-p53*KO cell line transplant prostate tumors. Scale bars, 50μm. **B**, Quantification of percentage of $CD31^+$ blood vessels with visible lumens from IHC staining in Fig. 4A (n = 3 mice per group). **C,** Representative flow cytometry gating strategy for expression of VEGFR2 and activation markers in spleen-derived $CD8^+$ T cells co-cultured with tumor cells *ex vivo* as in Fig. **4F-I**. **D,** Representative IHC staining in surgically resected primary prostate cancer patient samples stratified by MYC staining scores into high and low groups. Scale bars, 50μm. **E,** xCell analysis (55) of expression of transcripts of indicated $CD8^+$ T cell populations in primary prostate cancer patient samples from The Pan-Cancer Atlas stratified by expression of *KDR* (the gene encoding VEGFR2) into high and low groups. Data represent mean ± SEM. P-values were calculated by two-tailed, unpaired Student's t-test (**B**) and Wilcoxon test (**E**).
