## Supplementary Figure S3 for "MYC and p53 alterations cooperate through VEGF signaling to repress cytotoxic T cell and immunotherapy responses in prostate cancer"

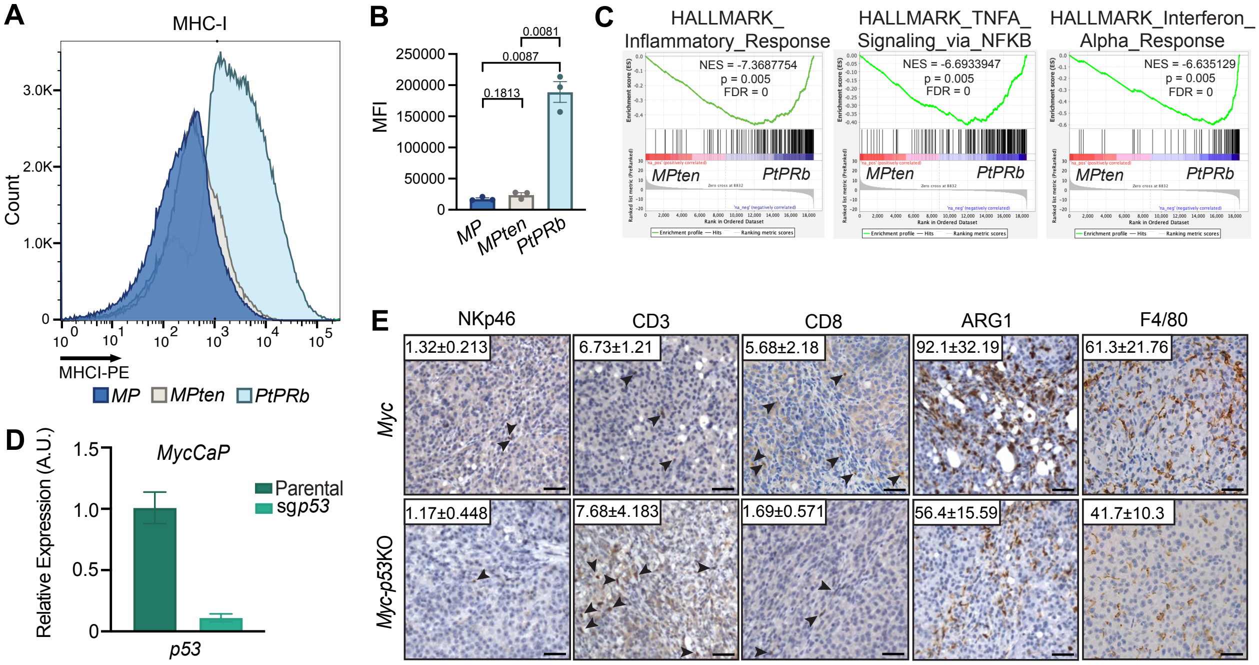


**Figure S3 (related to Figure 3). *p53* inactivation in MYC-driven prostate tumors leads to reduced MHC-I expression and CD8$^{+}$ T cell infiltration. A-B,** Representative histograms (**A**) and quantification (**B**) of mean fluorescent intensity (MFI) of MHC-I (H-2Kb) expression on *MP*, *MPten*, and *PtPRb* tumor cells (n = 3 biological replicates per group). **C,** Gene Set Enrichment Analysis (GSEA) of inflammatory, NF$\kappa$B, and interferon (IFN) signaling gene sets in indicated EPO-GEMM tumors (n = 8-17 mice per group). NES, normalized Enrichment Score. **D,** RT-qPCR analysis of *p53* expression in parental *Myc-CaP* and *Myc-p53*KO cells (n = 2 biological replicates associated with 3 technical replicates per group). **E,** Representative IHC staining of *Myc-CaP* (*Myc*) and *Myc-p53*KO cell line orthotopic transplant tumors in FVB mice harvested at endpoint. Arrowheads indicate positive staining for immune cells. Scale bars, 50µm. Quantification of NKp46$^{+}$ NK cells, CD3$^{+}$ and CD8$^{+}$ T cells, Arginase1$^{+}$ (ARG1) suppressive myeloid cells, and F4/80$^{+}$ macrophages per field is shown inset (n = 3 mice per group). Data represent mean ± SEM. P-values were calculated by two-tailed, unpaired Student's t-test.
