## Supplementary Figure S2 for "MYC and p53 alterations cooperate through VEGF signaling to repress cytotoxic T cell and immunotherapy responses in prostate cancer"

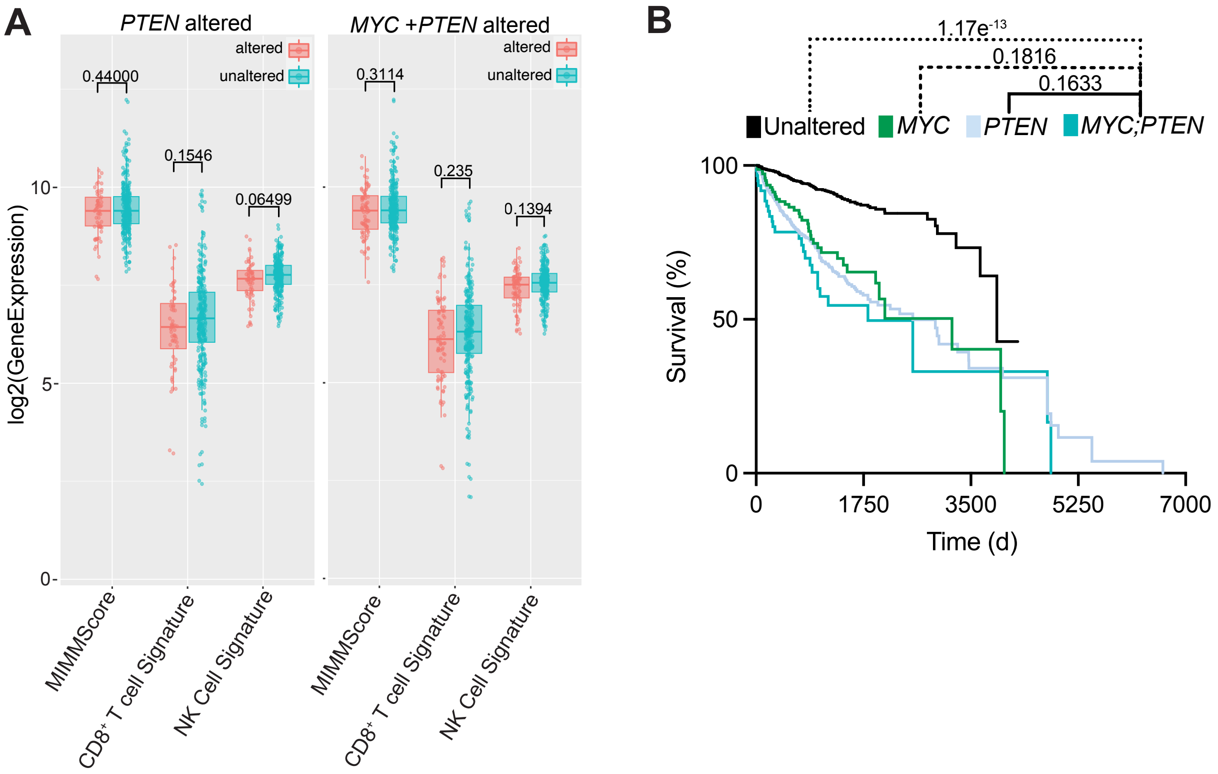


**Figure S2 (related to Figure 2). *MYC* and *PTEN* co-alterations do not cooperate to further drive immune suppression or lethality in human prostate cancer. A,** Box and whisker plots showing gene expression analysis of magnitude of immune infiltration [MImmScore (43)], NK cell (45), and CD8$^+$ T cell (44) signatures in primary prostate cancer patient samples from The Cancer Genome Atlas (TCGA) (42) stratified by alterations in *MYC*, *PTEN*, or compound *MYC;PTEN* (n = 79-235 per group). **B,** Kaplan-Meier survival curve of metastatic CRPC patients harboring prostate tumors with *MYC* or *PTEN* alterations alone or in combination from a SU2C dataset (15) (n = 17-143 per group). Data represent mean ± SEM. P-values were calculated by Wilcoxon test (**A**) and log-rank test (**B**).
